# Evolutionary reconstruction of bat influenza A viruses reveals segment-specific histories and host-associated selection

**DOI:** 10.64898/2026.10.14.754429

**Authors:** Austen Rowell, Matthew G. Johnson, David A. Rasmussen, Lisa M. Bono

## Abstract

Influenza A (IAV) is a major zoonotic pathogen infecting a broad range of vertebrate hosts. In this study, we focus on inferring the evolutionary history of a subset of novel IAV, isolated from New World bats, which use a non-canonical host receptor and infer their relationship to sialic-acid-binding IAV. Little is known about when these novel subtypes emerged and how selection has acted upon them compared to sialic-acid-binding circulating strains. The current state of the field is split on whether bats are natural reservoirs for a distinct yet low abundance IAV lineage, or if influenza underwent rapid evolution after a recent spillover from a species within its standard host range. By analyzing segment-specific phylogenies, we hope to clarify the evolutionary history of bat IAV. Our objectives are to infer the date of divergence of non-canonical bat IAV from sialic-acid binding IAV, identify patterns of selection acting on bat IAV compared to representative samples of sialic-acid-binding IAV to identify potential signatures of genome-wide adaptation, and finally to reconstruct ancestral state transitions to infer which host taxa preceded spillover into bats. To accomplish this, we analyzed each IAV genome segment independently using least-squares molecular dating, Bayesian discrete-trait ancestral host reconstruction, and branch-site selection tests. Our results suggest that non-canonical bat IAV diverged from major IAV lineages 100-200 years ago, supporting a deeper evolutionary history rather than a very recent spillover event. However, discordance among genome segments in inferred divergence times, selective pressures, and ancestral hosts indicates that bat IAV experienced segment-specific evolutionary histories. Together, these findings suggest that non-canonical bat IAV represents a deeply divergent lineage with a complex evolutionary history involving segment-specific selection and host-transition dynamics.

## Introduction

Influenza A virus infects a broad range of avian and mammalian hosts, yet most sequences of IAV are associated with birds, humans, and swine (Bao et al., 2008; Wolf et al., 2006). IAV has rarely been identified in bats, even though IAV can infect a variety of avian and mammalian species and bats often come into close contact with these species (Mikula et al., 2016). Despite this potential for exposure, IAV in bats remains rarely detected and poorly represented in sequencing databases potentially due to low screening in bats. This limited sampling makes it difficult to determine whether bat-associated IAV is poorly represented because of genuinely infrequent circulation, geographically or taxonomically restricted host associations, or technical and surveillance factors such as PCR primer mismatch and limited sampling of bats.

Despite the lack of sampling, the bat-associated IAVs detected to date include lineages with highly unusual and unique phenotypes, particularly among New World H17/H18 viruses. IAV was first isolated from bats in the New World. Despite having a high degree of sequence similarity with other IAVs, there are critical differences in the hemagglutinin (HA) and neuraminidase (NA) gene products, which affect attachment (and entry into) and cleavage (and exit from) the host cell. Structural and functional analyses showed that H17 and H18 do not attach to either alpha 2,3- or alpha 2,6-linked sialic acid, the canonical receptor for IAV (Li et al., 2012; Sun et al., 2013; Wu et al., 2014), but rather with major histocompatibility complex II proteins (MHC-II) (Ciminski et al., 2020; Karakus et al., 2020).

H17/H18 influenzas exhibit a broad MHC-II specificity, binding to MHC-II expressed in bats, mice, pigs, and chickens (Karakus et al., 2019). MHC-II-associated entry is not restricted to New World bat IAV. Novel H19 subtypes isolated from birds also bind to MHC-II, but H19 has a narrower host range (Karakus, Mena, et al., 2024). These findings have led to the discovery that H3N2 and H2N2 can enter cells using MHC-II independent of sialic acids (Cardenas et al., 2026; Karakus, Sempere Borau, et al., 2024; Hargadon et al., 2011).

Furthermore, unlike all other known IAVs, N10 and N11 are not sialidases, meaning that they cannot cleave from sialic acids during exit from the cell. These proteins have no known catalytic activity (Li et al., 2012). Although N12 has been less studied than N11, sequence differences between N11 and N12 suggest that there may be morphological difference between the two that may influence its function during the release of viruses (Echeverri-De la Hoz et al., 2025). These findings suggest that noncanonical receptor use is not an isolated feature of H17/H18, but the selective pressures and ecological interactions that generate and maintain these phenotypes remain poorly understood.

Previous phylogenetic studies established that bat-associated IAVs are divergent from canonical IAV lineages using single gene (HA) maximum likelihood phylogenies based on a subset of IAV sequence diversity (Ciminski et al., 2020; Tong et al., 2012). As HA is involved in attachment this is useful to identify distance based on host range and changes in tropism. However, the deep placement of bat-associated IAVs could reflect recent spillover from a non-bat host followed by rapid genome-wide adaptation to a new host environment, or alternatively, older and rarely sampled IAV diversity that has persisted in bats or related ecological niches over longer evolutionary timescales (He et al., 2019; Hill et al., 2022). Distinguishing between these models requires more than estimating relatedness among sequences and within a single gene tree (Lemey et al., 2009; Nabhan & Sarkar, 2012; Yang & Bielawski, 2000; Zwickl & Hillis, 2002). We therefore use a combination of divergence dating to test whether bat-associated IAVs are recently derived from sampled canonical IAV diversity or represent deeper lineages; selection analyses to evaluate whether adaptive signals are genome-wide or localized to specific branches and sites; and Bayesian ancestral host-state reconstruction to identify host states inferred upstream of bat-associated lineages. This integrative framework allows us to evaluate whether the unusual biology of bat-associated IAV reflects rapid adaptation following recent host jumps or a longer, more complex evolutionary history obscured by sparse sampling.

## Results

### Molecular dating supports older divergence of bat-associated IAV lineages

We first estimated segment-specific divergence times to distinguish recent spillover from deeper evolutionary separation. We inferred the time to most recent common ancestor (tMRCA) of novel bat influenza using maximum likelihood phylogenies combined with LSD2 molecular dating in IQ-TREE assuming a constant molecular clock rate (Minh et al., 2020). The tMRCA was determined at the parental node to the monophyletic bat IAV clade. This offers an estimate of how far diverged the bat IAV clade is from the rest of the samples in each gene tree. Due to computational complexity associated with the full dataset, we resampled IAV genomes to a representative set of 75 sequences using a phylogenetically informed approach. First, we retained the 25 sequences nearest to the H17/H18 clades to preserve close evolutionary context around the focal bat-associated lineages. The remaining sequences were selected using patristic distance to sample representatives from major clades across the broader tree. In *Figure 1*, we illustrate the variability in tMRCA found across IAV genes. Parental nodes of the bat clade from trees with >90 ultrafast bootstrap support values suggest a divergence estimate within the last 100 to 200 years (*Supplemental Table 1*). Expanded and 75-taxon subset tMRCA estimates were strongly correlated, indicating broadly consistent temporal patterns across segments, but the absolute date of the tMRCA estimates remain sensitive to taxon sampling (Pearson *r* = 0.894; Spearman *rho* = 0.905) (*Figure 2*). Overall, our analysis dates bat-associated IAV lineages centuries prior to their collection. We therefore next explore the selection pressures experienced by each gene to determine whether it is likely that these lineages underwent rapid evolution after a recent spillover or if their selective profiles suggest a long-term evolutionary history consistent with a lowly-abundant yet persistent population within bats.

**Figure 1:**
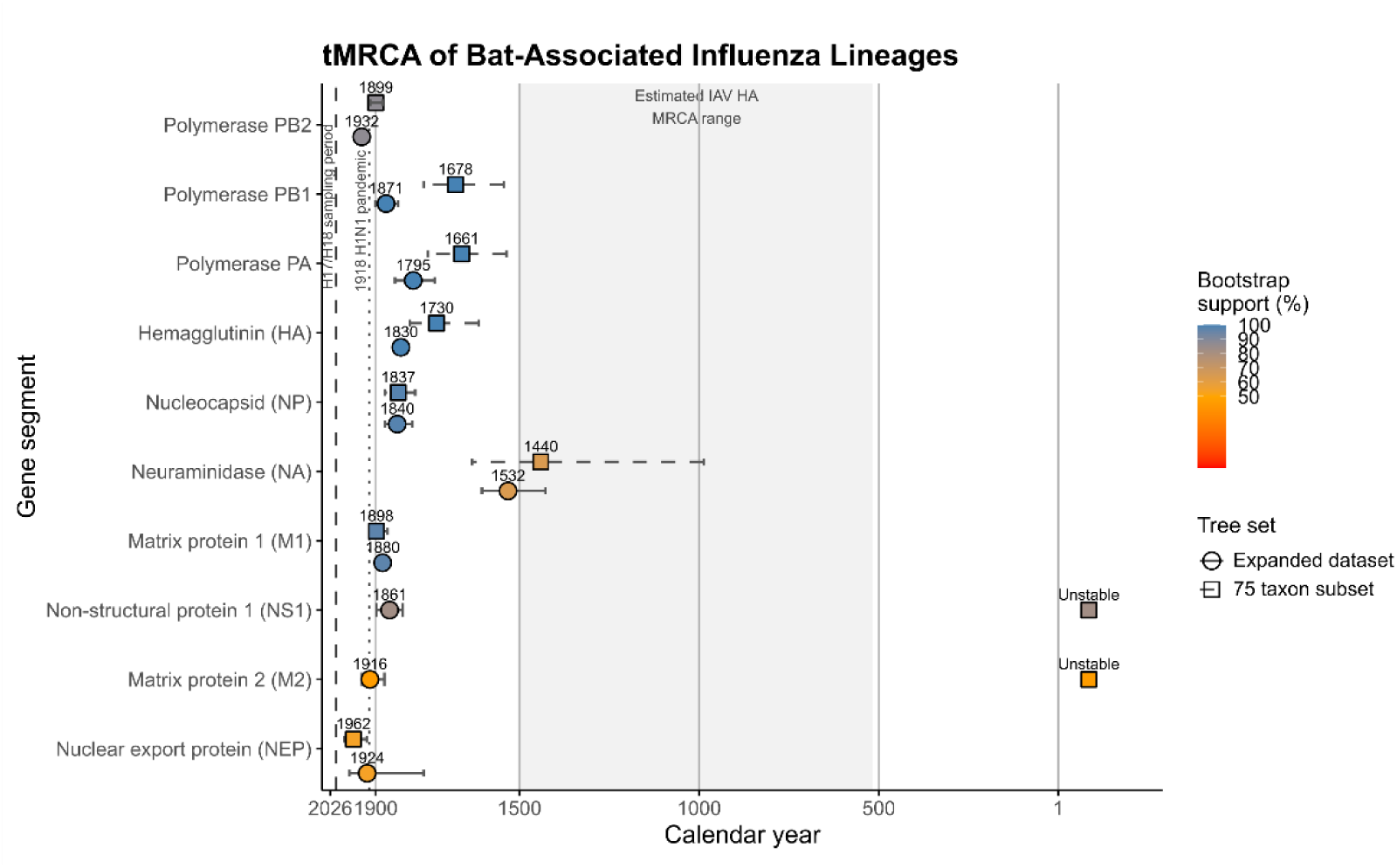
Estimated tMRCAs for novel bat IAV gene segments. Points represent median LSD2 tMRCA estimates, with horizontal bars showing confidence intervals derived from 1,000 LSD2 replicates. Point color indicates ultrafast bootstrap support for the internal node representing each segment-specific MRCA, with blue indicating higher support and red indicating lower support. The x-axis is scaled by distance from 2026 and labeled by calendar year. Vertical reference lines indicate selected IAV events, including the bat IAV H17/H18 sampling period and the 1918 H1N1 and 1968 H3N2 pandemic emergence events. The shaded region indicates a published estimated MRCA range for influenza A hemagglutinin proteins and is included as a broad evolutionary reference.

**Figure 1:**
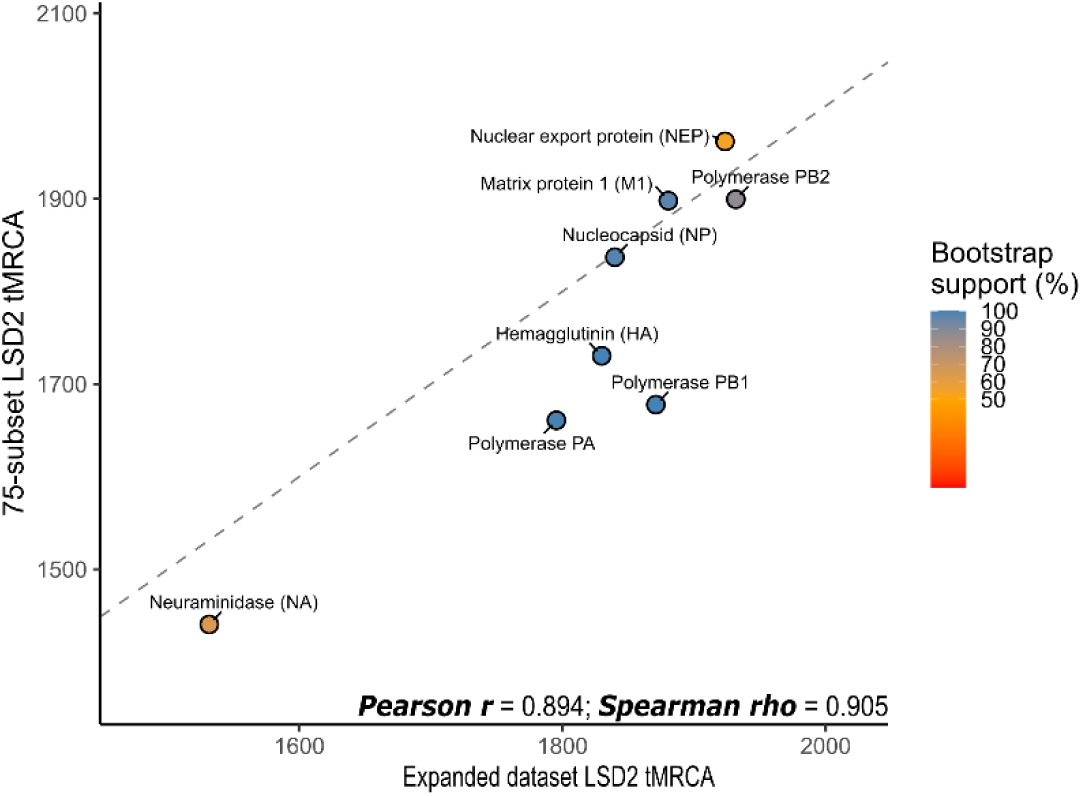
Comparison of LSD2 tMRCA estimates inferred from the expanded dataset and 75-taxon subset. Each point represents a bat-associated MRCA estimate for one influenza gene segment, colored by UFBoot support for the corresponding MRCA node or equivalent split. The dashed line indicates a 1:1 relationship between expanded and subset estimates. Expanded and subset tMRCA estimates were strongly correlated after excluding unstable NS1 and M2 subset estimates, although several genes showed shifts in absolute date estimates.

### Little evidence for a shift in the strength of selection in bat IAV

The strength of selection experienced by a specific site or branch of a tree can be used to infer how changes in the environment influence the directionality of evolution within a lineage. We used both site and branch level codon substitution models to determine whether there is atypically strong or weak selective pressures acting on particular branches, such as our bat IAV clade, or specific amino acid residues. In this case as the viral environment is often considered as the host they are infecting, we can infer how changes in host may have impacted the evolution of bat IAV assuming bat-associated IAV arose through host transition from a non-bat lineage. First, we tested if the strength of selection has intensified or relaxed along our branch of interest using RELAX (Wertheim et al., 2015). RELAX determines the distribution of ω=dN/dS across sites within foreground (i.e. bat) and reference (non-bat) branches. It then introduces a selection intensity parameter (K) that transforms the ω values on the foreground branch. Specifically, foreground ω values are modeled as ωiK, such that K>1 indicates an intensification of selection on the foreground branch and K<1 indicates a relaxation in selection—this is not associated with directionality for purifying or diversifying selection, but instead a change in the overall distribution of ω values towards or away from ω=1. A result of K=1 indicates there is no detectable shift in the strength of selection experienced along the foreground branch.

Using the same subset used for the dating analysis reduced overrepresentation of densely sampled clades with near-identical sequences while maintaining phylogenetic diversity across the dataset. We used the representative set of 75 sequences for each gene to test for a shift in strength of selection. RELAX analysis of bat IAV branches showed evidence for intensification of selection in only one gene (M2), while the rest showed either no shift or relaxation of the magnitude of selection along bat IAV branches (*Table 1*).

**Table 1:** Selection analysis summary for bat-associated influenza gene segments. RELAX identifies shifts in selection intensity, while PAML branch-site, aBSREL, BUSTED, MEME, and FUBAR test for evidence of episodic or pervasive selection. RELAX estimates a selection intensity parameter (K), where K > 1 indicates intensified selection and K < 1 indicates relaxed selection. Bold values indicate statistically significant results. FUBAR site counts represent sites with posterior probability ≥0.90.

| Gene | RELAX |  |  | PAML branch-site |  | aBSREL |  | BUSTED |  | MEME | FUBAR |
| --- | --- | --- | --- | --- | --- | --- | --- | --- | --- | --- | --- |
| | K | p-value | Direction | LRT | p-value | LRT | q-value | LRT | p-value | Sites with $p \leq 0.05$ | Sites with PP $\geq 0.90$ |
| Polymerase PB2 | 1.031 | 0.744 | No shift | 193.711 | **2.46E-44** | 15.823 | **1.24E-04** | 1.061 | 0.294 | 8 | 1 positive; 734 purifying |
| Polymerase PB1 | 0.947 | 0.714 | No shift | 56.622 | **2.64E-14** | 0 | 1 | 5.499 | **0.032** | 18 | 0 positive; 721 purifying |
| Polymerase PA | 0.639 | **6.11E-15** | Relaxed | 62.703 | **1.20E-15** | 8.648 | **0.005** | 6.887 | **0.016** | 11 | 1 positive; 673 purifying |
| Hemagglutinin (HA) | 0.999 | 0.978 | No shift | 10.439 | **6.17E-04** | 0.477 | 0.334 | 0 | 0.5 | 8 | 1 positive; 572 purifying |
| Nucleocapsid (NP) | 0.687 | **2.52E-04** | Relaxed | 46.509 | **4.56E-12** | 42.987 | **1.50E-10** | 48.091 | **1.80E-11** | 0 | 0 positive; 460 purifying |
| Neuraminidase (NA) | 0.997 | 0.955 | No shift | 25.487 | **2.23E-07** | 0 | 1 | 0.012 | 0.497 | 4 | 0 positive; 473 purifying |
| Matrix protein 1 (M1) | 1 | 1 | No shift | 8.784 | **0.002** | 6.116 | **0.017** | 1.02 | 0.3 | 18 | 0 positive; 219 purifying |
| Matrix protein 2 (M2) | 8.404 | **0.045** | Intensified | 12.757 | **1.77E-04** | 3.723 | 0.057 | -0.006 | 0.5 | 24 | 10 positive; 35 purifying |
| Nonstructural protein 1 (NS1) | 0.523 | 0.0117 | Relaxed | 23.954 | **4.93E-07** | 23.185 | **3.06E-06** | 23.779 | **3.43E-06** | 0 | 19 positive; 146 purifying |
| Nuclear export protein (NEP) | 1 | 1 | No shift | 9.998 | **7.83E-04** | 0 | 1 | 0 | 0.5 | 8 | 2 positive; 126 purifying |

### Evidence for episodic selection in bat IAV

Branch- and site-level selection analyses were used to test whether bat-associated influenza lineages showed evidence of episodic or directional selection. In viruses with broad host ranges or histories of host switching, such signals can indicate adaptive changes associated with host-specific selective pressures. Here, we used PAML branch-site analysis, and complementary HyPhy methods, including aBSREL, BUSTED, MEME, FUBAR, and FADE to evaluate evidence for selection across influenza gene segments (*Table 1; Supplemental Table 2*) (Yang, 1997; Pond et al., 2005). PAML branch-site analysis was used to test for episodic positive selection along the bat-associated foreground lineage. This model compares a null model, in which sites in the foreground lineage are constrained to ω = 1, against an alternative model that allows ω > 1 at a subset of codons along the foreground lineage. A likelihood-ratio test was then used to determine whether the alternative model provided a significantly better fit. Multiple genes showed significant support for the branch-site alternative model allowing for ω > 1, indicating that positive selection was consistent with a subset of codon sites along the foreground lineage, rather than a uniform elevation of ω across the entire gene. Site-level methods showed additional evidence that selection was concentrated at specific residues rather than uniformly distributed across entire genes. Bayes Empirical Bayes analysis identified candidate positively selected sites within most genes except for NS2 and PB2, with sites exceeding posterior probability thresholds of 0.95 or 0.99 (*Figure 3 & Supplemental Table 2*). The highest concentration of sites were found in PB1 (15), NP (12), and NA (44).

**Figure 3:**
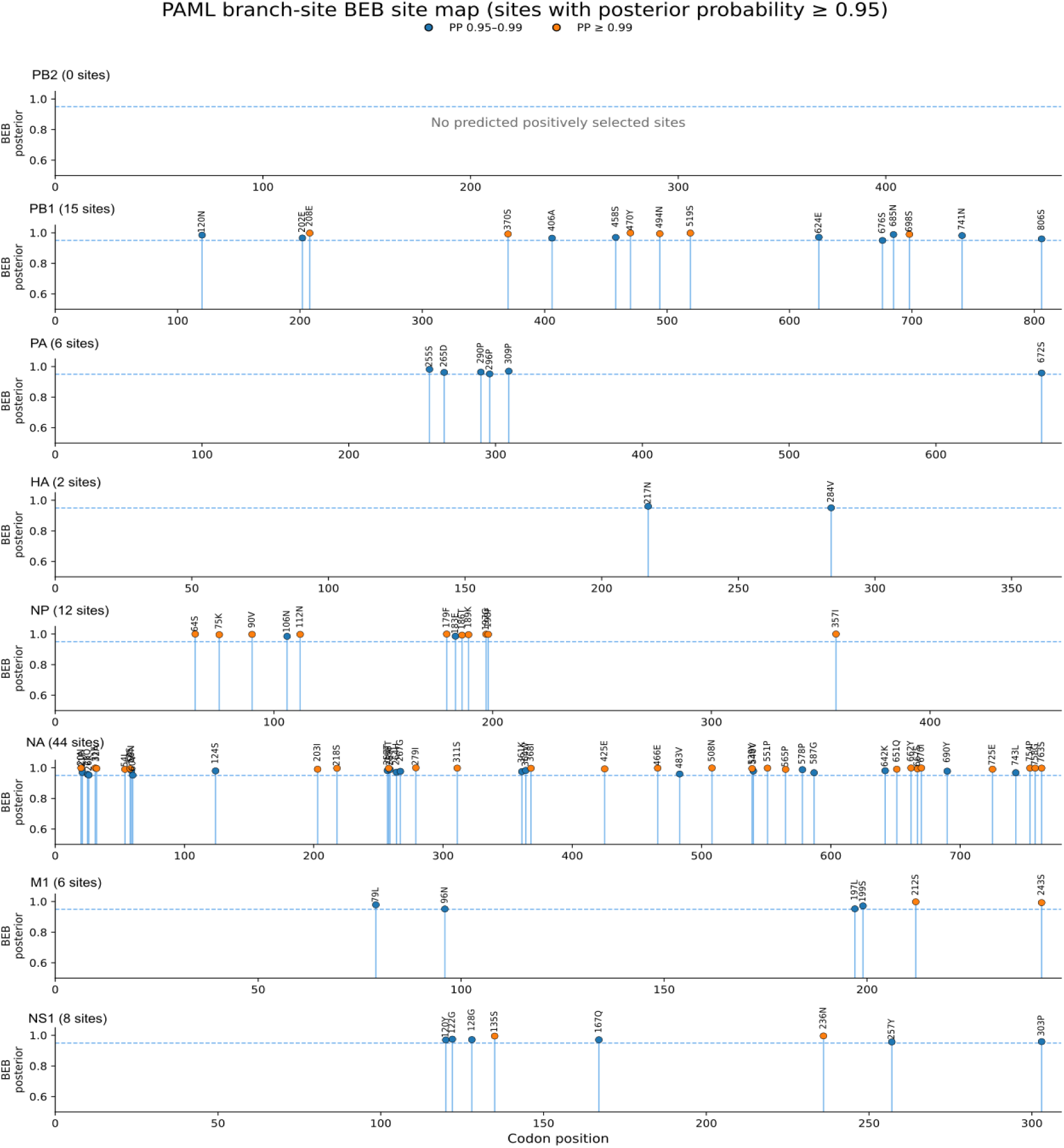
Distribution of PAML branch-site BEB-inferred positively selected sites across influenza gene segments. Points indicate sites with BEB posterior probability ≥0.95; orange points indicate sites with posterior probability ≥0.99. Codon positions correspond to PAML-reported codon positions from the codeml branch-site output. Exact site identities and posterior probabilities are provided in Supplementary Table X.

Similarly, aBSREL detected evidence of episodic diversifying selection on bat IAV foreground branches for PB2, PA, NP, M1, and NS1. BUSTED tested for gene-wide evidence that at least one site experienced episodic diversifying selection on the foreground lineage. These tests are related but ask different questions which explain differences in significance between methods. For example, a gene may be significant in BUSTED if selection occurred at a subset of sites anywhere on the foreground lineage but may not be significant in aBSREL if the signal is not concentrated on a particular branch.

MEME tests for episodic selection at individual codon sites, whereas FUBAR detects pervasive selection across the phylogenetic tree. As a result, MEME and FUBAR are expected to differ when selection is transient or lineage-specific such as we find in bat IAV. Sites detected by MEME may represent episodic selection acting on particular branches within the phylogeny, whereas FUBAR is more likely to identify sites under consistent diversifying or purifying selection across the tree. FADE further complements these analyses by testing for directional amino acid preferences rather than shifts in nonsynonymous substitution rates alone. FADE did not find evidence for a directional shift in amino acid preference in any gene tree, suggesting that noncanonical receptor use may be an under-recognized feature of IAV evolution. In this dataset MEME identified relatively few sites with evidence of diversifying selection, whereas FUBAR identified most significant sites as being under purifying selection. Cross-method comparison identified 35 gene-site combinations supported by at least two positive-selection methods (*Figure 4*). Most overlapping sites were detected by both MEME and FUBAR, indicating agreement between site-level methods despite their different assumptions. Only six sites were supported by PAML branch-site analysis and at least one additional method, suggesting that relatively few site-level signals overlapped with evidence for positive selection on the bat-associated foreground lineage. This pattern indicates that positive-selection signals were concentrated in a limited number of residues, while most codons remained evolutionarily constrained. Sites supported by both PAML and a complementary site-level method represent the strongest candidates for bat-lineage-associated positive selection.

**Figure 4:**
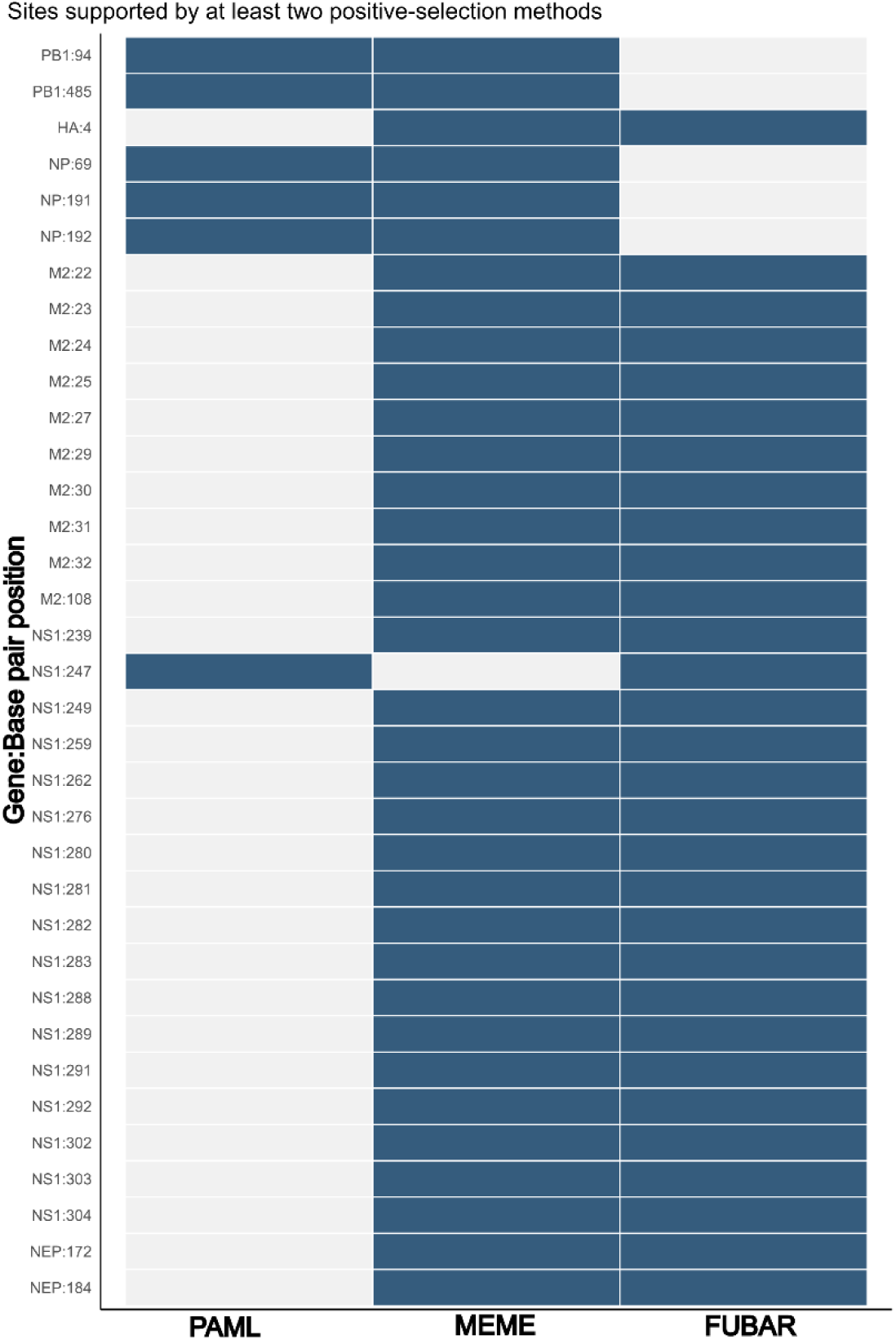
Cross-method agreement among sites inferred to be under positive selection. The heatmap shows 35 gene–site combinations that met the selection criterion for at least two methods. Filled cells indicate support from PAML branch-site Bayes empirical Bayes analysis (posterior probability ≥0.95), MEME (nominal p<0.05), or FUBAR (posterior probability ≥0.90); unfilled cells indicate that the site was not identified as significant.

### Codon-usage profiles differ in bat IAV

To determine whether the detected patterns of widespread episodic selection could arise from changes in codon-usage or bias we chose the metrics of relative synonymous codon usage (RSCU), overall GC content, GC third base pair content (GC3), or effective number of codons (ENC) to determine whether there are specific amino acid residue shifts, or if there are overall profile shifts within bat IAV lineages compared to the rest of the phylogeny. PERMANOVA analysis of RSCU profiles identified significant host-associated codon usage differences for multiple gene segments, including PB2, PB1, PA, M2, and NS1 after Benjamini–Hochberg (BH) correction for multiple testing. The strongest PERMANOVA effect sizes were observed for M2, PA, PB2, PB1, and NS1, although the proportion of explained variance remained modest across genes. Dispersion testing was used to determine whether these PERMANOVA signals could be explained by differences in within-group variability. The genes showed non-significant dispersion tests, suggesting that their PERMANOVA results likely reflect differences in host-associated codon usage composition (*Table 2*).

**Table 2:** RSCU profiles significantly differ in bat IAV lineages. PERMANOVA analysis of host-associated codon usage differences across influenza gene segments. Relative synonymous codon usage profiles were compared among host groups for each gene segment using PERMANOVA with 999 permutations. Several segments, including PB2, PB1, PA, NA, M2, and NS1, showed significant host-associated differences after Benjamini–Hochberg correction. Effect sizes were modest across genes, indicating that host group explains a detectable but limited proportion of codon usage variation.

| Gene | Permanova |  |  |  |  |
| --- | --- | --- | --- | --- | --- |
|  | sumsq model | sumsq residual | r2 | pseudo F | q value BH |
| Polymerase PB2 | 290.97 | 4075.03 | 0.07 | 5.21 | 2.50E-03 |
| Polymerase PB1 | 272.09 | 4093.91 | 0.06 | 4.85 | 4.00E-03 |
| Polymerase PA | 316.97 | 4049.03 | 0.07 | 5.71 | 2.50E-03 |
| Hemagglutinin (HA) | 68.34 | 4297.66 | 0.02 | 1.16 | 1.33E-01 |
| Nucleocapsid (NP) | 139.75 | 4226.25 | 0.03 | 2.41 | 6.43E-02 |
| Neuraminidase (NA) | 94.80 | 4271.20 | 0.02 | 1.62 | 7.33E-02 |
| Matrix Protein 1 (M1) | 111.64 | 4254.36 | 0.03 | 1.92 | 7.33E-02 |
| Matrix Protein 2 (M2) | 345.94 | 4020.06 | 0.08 | 6.28 | 2.50E-03 |
| Non-structural Protein 1 (NS1) | 221.39 | 4144.61 | 0.05 | 3.90 | 2.50E-03 |
| Nuclear Export Protein (NEP) | 151.82 | 3388.18 | 0.04 | 2.64 | 6.43E-02 |

To identify which codon-bias features contributed to host-associated differences, we compared individual codon-usage metrics between bat and non-bat sequences using Wilcoxon rank-sum tests with BH correction. Several significant differences were detected in GC-related metrics. Bat-associated sequences showed lower GC-related values for multiple genes, including M2, PA, PB2, and M1. These results suggest that host-associated RSCU differences may partly reflect shifts in nucleotide composition at synonymous sites, particularly GC3 and total GC content. However, not all genes with significant PERMANOVA results showed significant univariate metric differences, indicating that host-associated codon usage variation may also be distributed across multiple codons rather than captured by a single summary statistic (*Supplemental Table 3*). These results indicate that bat-associated sequences differ modestly in codon-usage composition, with some differences associated with GC-related metrics. However, because codon usage can reflect both mutational processes and host-associated selection, these patterns should be interpreted as compositional differences rather than direct evidence of adaptive codon optimization.

### Ancestral host reconstruction reveals discordant or unresolved histories

An outstanding question for these novel lineages is whether their genomic features and evolutionary histories indicate prior circulation in avian or mammalian hosts. The long-standing view in influenza ecology is that wild birds, particularly waterfowl, represent the major natural reservoir of influenza A virus diversity. Therefore, we asked whether ancestral host reconstruction could identify the most likely host state preceding the bat-associated IAV clade and whether this history implicated prior evolution in avian hosts, humans, or other mammals. To test this, we used a continuous-time, discrete-trait phylogeographic model to estimate the posterior distribution over ancestral host states across each gene tree. Host metadata were encoded into five broad host states to reduce model complexity and limit the influence of uneven sampling among host groups. Because discrete-trait reconstructions are sensitive to sampling bias and prone to high levels of uncertainty regarding ancestral states deep in the past, these results should be interpreted as probabilistic reconstructions of host state history rather than direct evidence of host jumps (Layan et al., 2023; Maio et al., 2015).

We used an asymmetric discrete-trait model with Bayesian stochastic search variable selection to estimate host-transition rates (Lemey et al., 2009). This approach allows transition rates between host states to vary while identifying the subset of rates that are most strongly supported by the tree and trait data. Posterior Markov-jump histories were summarized along the path from the root to the MRCA of the H17/H18-associated lineage for each gene segment (*Figure 5A*). Most interpretable segments reconstructed the MRCA as bat, but the nearest upstream non-bat host varied among genes with reconstructions showing a high degree of uncertainty. PB2 showed the strongest upstream-source signal, with avian inferred as the nearest non-bat state in 82.5% of posterior histories, while HA also most often supported an avian upstream state in 50% of posterior histories. Other segments inferred human, environment, or non-human mammal as the most common upstream state, but posterior support was low, indicating that the exact source state could not be confidently resolved from the sampled data (*Figure 5B*). These results support a model in which New World bat IAVs are deeply divergent from sampled canonical IAV diversity, but their genome segments do not share a single uniform history. Instead, divergence times, selection signals, codon-usage profiles, and inferred ancestral host states vary among segments, consistent with a complex evolutionary history potentially shaped by reassortment, host-associated constraints, and sparse sampling of bat IAV diversity.

**Figure 5:**
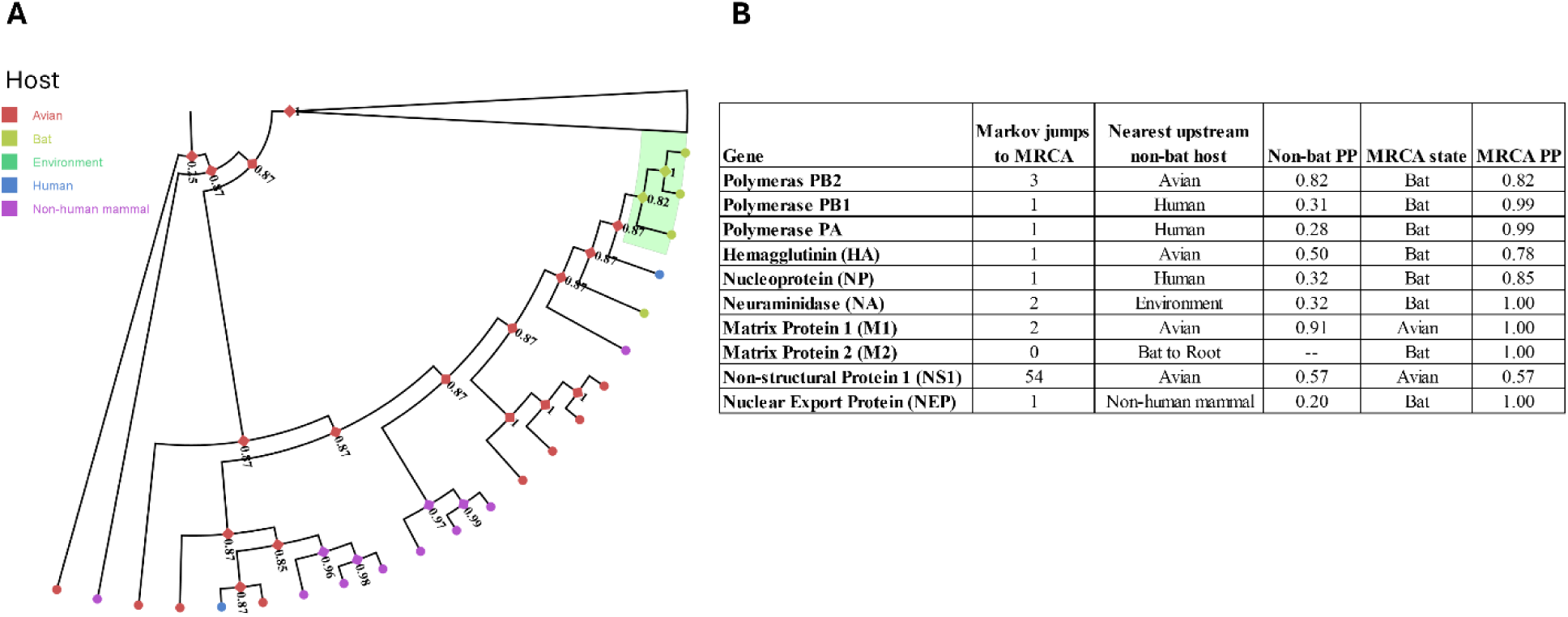
Ancestral host-state reconstruction across bat influenza gene segments. **(A)** Cladogram of discrete-trait history tree for the polymerase PB2 segment illustrating the host-reconstruction framework applied across gene trees. Tip colors indicate observed host and internal nodes are colored by inferred host category, and the highlighted lineage marks the target bat influenza clade. Numeric node labels indicate posterior probability support for the displayed ancestral-state reconstruction. **(B)** Summary of posterior Markov-jump histories from the root to the MRCA of the focal H17/H18-associated lineage for each gene segment. The table reports median inferred host-state transitions, the nearest upstream non-bat host state when resolved, posterior support for that state, and the inferred host state at the focal MRCA. Additional gene trees are provided in the supplement.

## Discussion

The evolutionary origin of H17/H18 bat-associated IAV remains difficult to resolve because these lineages are deeply divergent, sparsely sampled, and have segment-specific histories. By integrating molecular dating, selection analyses, codon-usage comparisons, and ancestral host-state reconstruction, we evaluated whether these viruses are more consistent with recent host transition followed by rapid genome-wide adaptation or with older, under-sampled viral diversity shaped by segment-specific evolution. Our results favor the latter model: bat-associated IAV lineages appear to represent older, under-sampled viral diversity rather than a strictly recent spillover followed by rapid genome-wide adaptation. Deep tMRCA estimates suggest long evolutionary timescales, while RELAX provides limited evidence for widespread intensification of selection. Instead, significant branch- and site-level signals from PAML branch-site, aBSREL, BUSTED, and FUBAR suggest episodic positive selection acting on specific lineages or residues, consistent with localized adaptation over a longer evolutionary timescale.

Reassortment likely explains the discordant segment-specific evolutionary histories of bat IAV. IAV, as a segmented virus, has the capacity to reassort during superinfection (Hockman et al., 2020). This swapping generates diversity and novelty for viral populations. Instances of reassortment have been linked to zoonoses or host range expansion events (Ganti et al., 2022).

Our phylogenetic and selection analyses support a long evolutionary history of adaptation within bats as opposed to a recent spillover. First, RELAX did not find evidence for a shift in selection across most genes, but branch-site methods did support sites under diversifying selection in bat lineages against a background of purifying selection acting on most sites in each gene. These results are more consistent with localized, lineage- or site-specific adaptation than with rapid genome-wide adaptation. Second, we found support for long-term localized adaptation from the codon usage analyses. We compared how sequence composition differed in the MHC-II binding bat IAV and sialic-acid-binding IAV and found modest but significant differences in host-associated compositional metrics in M2, PA, PB2, PB1, and NS1. These significant differences in RSCU and GC content could be the result of host adaptation, coinciding with changes to a few codon sites which may be advantageous within a particular host.

Viruses are obligate intracellular pathogens that rely on host cellular machinery for replication, transcription, translation, and immune evasion. As a result, viruses are expected to experience host-associated selective pressures, although these pressures may be unevenly distributed across genes and sites rather than producing genome-wide adaptive change. This pattern is consistent with our results: selection signals and codon usage were not distributed uniformly across the genome, but were concentrated in a subset of genes and residues with known roles in host interaction. The polymerase subunits PA, PB2, and PB1 are frequently associated with host-specific adaptation because of their roles in cap-snatching and replication (Domingues et al., 2019). PB1 has been implicated in degradation of mitochondrial antiviral signaling protein (MAVS), which can reduce interferon signaling (Rashid et al., 2023), while NS1 is a major innate immune antagonist that suppresses host interferon responses (Tisoncik et al., 2011). This is relevant because interferon activation in bats differs markedly from other mammals, and bat-associated IAV have been shown to have altered function (Chauché et al., 2018; Glennon et al., 2015; Irving et al., 2021; Turkington et al., 2018; Xie et al., 2018). These patterns do not demonstrate that bat IAV evolved directly in response to interferon activation, but they are consistent with the possibility that host immune environment contributed to gene-specific selective pressures.

Spillover or transmission from one host to another is dependent on overcoming ecological, cell-entry, and immunological barriers. To explore host jumps that may have potentiated successful spillover into bats, we used ancestral host-state reconstruction to identify host states upstream of the bat-associated clade. The reconstructions support spillovers into bats from multiple hosts rather than a particular type of host. PB2 provided the strongest support for an avian upstream state, and HA showed weaker avian support. Other segments had low posterior support for human, environmental, or non-human mammalian upstream states. In several cases, the host-reconstruction results did not identify a clear non-bat host immediately preceding the bat-associated lineage. For some segments, bats were reconstructed as the ancestral host at or near the root of the tree, leaving no well-supported upstream non-bat host. For other segments, the MRCA of the focal H17/H18 accessions also included non-bat or distantly related lineages, meaning that the reconstructed MRCA did not represent a restricted bat-associated clade alone. These cases limited our ability to infer a specific host state preceding the bat-associated lineage. Thus, the host-reconstruction results support segment-specific and often unresolved host-state histories rather than a single, well-resolved host transition into bats.

Historical opinion in the field is that avian species, particularly waterfowl, act as the natural reservoir for IAV (Diskin et al., 2020). A common cell-entry barrier for avian-to-mammal spillover is that avian-adapted IAV preferentially use α2,3 sialic acid receptors whereas mammalian-adapted IAV more often use α2,6 sialic acid receptors (Eriksson et al., 2018; Ferreri et al., 2022). Therefore, spillover may be facilitated through an intermediary species which has α2,3 sialic acid and α2,6 sialic acid co-localized providing an opportunity for IAV variants to access either cellular receptor (Parrish et al., 2015; Webby et al., 2000). Thus, if avian or other non-bat hosts contributed to the ancestry of these lineages, the transition into bats may have involved an intermediary host, bat species with permissive receptor distributions, or historical use of non-canonical receptor pathways. Consistent with the possibility of permissive receptor distributions, certain bat species (*Myotis lucifugus*) have co-localized sialic acid receptors allowing avian- and mammalian-adapted IAV infections in their gastrointestinal (GI) tract (Chothe et al., 2017). In addition, the GI tract has been shown to have gut-associated lymphoid tissue which possess MHC-II receptors which can permit H18N11 infection (Kessler et al., 2024). However, the tissue-specific distribution and co-localization of sialic acid receptors and MHC-II are not well characterized across bat species. As a result, it remains difficult to determine whether historical host transitions occurred directly within bats, involved intermediary hosts, or occurred after non-canonical receptor use had already evolved; all three possibilities remain plausible.

The primary limitation of this study is the sparse and uneven sampling of bat-associated IAV. This limitation was especially apparent in the Markov-jump summaries, where the bat-associated MRCA was often reconstructed with higher confidence than the identity of the nearest upstream non-bat state. Current IAV sequence databases are heavily shaped by outbreak surveillance, agricultural sampling, and human health priorities, leading to dense sampling of some hosts, time periods, and geographic regions but limited representation of wildlife reservoirs. This imbalance is most likely to affect ancestral host reconstruction because inferred host states and transition histories depend directly on the distribution of sampled hosts across the tree. For molecular dating, sparse sampling would primarily affect MRCA estimates if earlier-diverging or unsampled bat-associated lineages alter the inferred depth of the bat clade. For selection and codon-usage analyses, the major limitation is whether currently available bat IAV sequences are representative of bat-associated IAV diversity as a whole. Our subsampling strategy was designed to retain phylogenetic diversity and reduce overrepresentation of densely sampled clades, but it cannot correct for biases already present in public databases. Broad host categories also reduced model complexity but may obscure finer-scale ecological differences among host species. Additional bat IAV genomes, broader sampling of potential intermediary hosts, and richer metadata would substantially improve the ability to distinguish between long-term bat circulation, repeated spillover, and reassortment-driven histories.

Overall, our analyses provide an evolutionary and ecological framework for understanding non-canonical bat IAV. Rather than supporting a simple model of recent spillover followed by rapid genome-wide adaptation, the results point to older divergence, segment-specific histories, and localized branch and site-specific selection pressures. Molecular and structural studies have been essential for identifying the unusual receptor biology of bat IAV, but evolutionary analyses provide complementary context for how such phenotypes may arise and persist. Continued surveillance of IAV in bats and other under-sampled wildlife hosts will be necessary to determine whether MHC-II associated IAV phenotypes are rare evolutionary outliers or part of a broader, under-recognized diversity of influenza host interactions.

## Materials and Methods

### Data Accession

We downloaded sequence data and associated metadata from NCBI Virus on 24 July 2025 (Brister et al., 2015). We filtered sequences by genome segment and curated the dataset to remove laboratory-passaged isolates, vaccine strains, and entries with <90% genome completeness. Codon alignments, metadata, scripts, and beast xml files are deposited under DOI: https://doi.org/10.5281/zenodo.23346033.

### Alignment

We performed codon-aware multiple sequence alignment using MACSE, with internal stop codons masked to enable downstream branch-site selection analyses (Ranwez et al., 2018). We removed near-identical sequences using 95% similarity centroid clustering (Rognes et al., 2016). We inferred an initial phylogeny with FastTree v2.1.11 and used it as a guide tree for PARNAS v0.1.7 subsampling, which retained taxa representing ≥98% of the observed sequence diversity while reducing the dataset to a tractable size for subsequent analyses (Price et al., 2009; Markin et al., 2023).

### Time-scaled Phylogenies

We inferred time-scaled phylogenies for each gene using IQ-TREE v2.4.0 with codon-aware alignments to predict when divergence of bat IAV may have occurred for each gene tree (Minh et al., 2020). Prior to molecular clock dating, we evaluated temporal signal using root-to-tip regression in TempEst by comparing sampling dates against root-to-tip genetic distances (Rambaut et al., 2016). We provided tip dates allowing calibration of branch lengths in units of time using the Least Square Dating (LSD2) method. We estimated trees under the GTR+F+I substitution model with 1000 ultrafast bootstrap replicates and approximate likelihood ratio tests to assess branch support. We used the resulting time-scaled trees to estimate the time to the most recent common ancestor (tMRCA) of the monophyletic bat clade for each gene, with confidence intervals derived from LSD2 replicates. We did not specify outgroups, as rooting was inferred from the IQ-TREE. Because temporal signal varied among gene segments and some trees contained long-branch outliers, tMRCA estimates were interpreted as approximate divergence-time estimates rather than definitive dates of host transition.

### Host transition analysis

To identify potential hosts that preceded the spillover into bats, we used a discrete-trait reconstruction model implemented in BEASTX v10.5 using a fixed tree topology inferred from IQ-TREE (Ayres et al., 2019; Baele et al., 2025; Minh et al., 2020). For host reconstruction, we encoded host metadata as a five-state discrete trait (human, non-human mammal, bat, avian, and environmental). We used these states to reduce model complexity and limit noise caused by rare host representation. We inferred ancestral host states and transition rates with a continuous-time Markov chain (CTMC) discrete-trait model using asymmetric transition rates with Bayesian stochastic search variable selection (BSVSS) (Lemey et al., 2009). We used a General Time Reversible (GTR) substitution model with gamma distributed rate variation across sites and an uncorrelated relaxed clock. We included collection dates derived from sequence metadata for temporal calibration (Tavaré, 1986; Yang, 1994; Edwards et al., 2011). We used a constant population size coalescent tree prior in lieu of detailed demographic information across multiple hosts. These parameters allowed branch lengths to vary while maintaining overall tree topology so that transition estimates were conditioned on tree topology and the observed tip host states.

### Selection analysis and branch site tests

We conducted selection analyses using PAML branch-site models and complementary HyPhy (v.2.5.8) methods. To create computationally tractable foreground-selection datasets, full segment alignments and trees were reduced to representative 75-sequence subsets using a phylogenetically aware approach (Markin et al., 2023). We retained all bat-associated accessions present in both the alignment and tree. We then removed non-bat sequences in which more than 50% of codon positions contained ambiguous or gap-containing codons. After filtering, we retained the 25 taxa with the smallest patristic distance based on branch length to any bat-associated tip, preserving close evolutionary context around the focal bat lineage. We selected the remaining background taxa using a farthest-first traversal across the tree to preserve broad phylogenetic diversity while avoiding overrepresentation of densely sampled clades. We used resulting sequences to create a new raw subset sequence file which was aligned in-frame by MACSE using a codon-aware method, and had a tree inferred using the same IQTREE parameters (Minh et al., 2020; Ranwez et al., 2018). For foreground-based selection analyses, test branches were defined as the bat-associated lineage containing H17N10 and H18N11 IAV sequences. Gene-specific trees were labeled at the bat-associated MRCA when the sequences formed a suitable monophyletic clade. Genes for which the foreground corresponded to the root or could not be unambiguously assigned were excluded from those specific analyses or treated separately as noted.

#### Changes in selection intensity: RELAX

To test for changes in the overall strength of selection associated with branches of interest, we used RELAX (Wertheim et al., 2015). This method estimates a selection intensity parameter (*K*), where *K* > 1 indicates intensified selection and *K* < 1 indicates relaxed selection on the test branches relative to the reference branches. Branches of interest were designated as those leading to and including H17N10 and H18N11 IAV to investigate the selective shifts associated with the bat-derived influenza lineages. RELAX analyses were conducted separately for each gene. Statistical significance was assessed using likelihood ratio tests, with *p* < 0.05 indicating a significant change in selection intensity.

#### Branch-specific episodic selection: aBSREL

Adaptive Branch-Site Random Effects Likelihood (aBSREL) was used to test for episodic diversifying selection acting on individual branches of the phylogeny (Smith et al., 2015). This method allows the nonsynonymous to synonymous substitution rate ratio (ω = dN/dS) to vary both among sites and across branches. Statistical significance was assessed using likelihood ratio tests (LRTs), with multiple-testing correction applied across branches within each tree using the Holm–Bonferroni procedure (Holm, 1979). Branches with corrected p-values below 0.05 were considered to show evidence of episodic diversifying selection.

#### Gene-wide episodic selection: BUSTED

To test for gene-wide episodic positive selection, we applied the Branch-site Unrestricted Statistical Test for Episodic Diversification (BUSTED) (Murrell et al., 2015). BUSTED was used to test whether any site experienced episodic diversifying selection on the bat-associated test branch set for each gene. BUSTED tests whether at least one site experienced episodic diversifying selection on the specified bat-associated test branch set. Although the model is fit using the full phylogeny, significance is interpreted as evidence for ω > 1 on the foreground branch set. Statistical significance was evaluated using a likelihood ratio test comparing the null model (ω constrained to ≤1) to the alternative model allowing ω > 1. A p-value < 0.05 was interpreted as evidence of episodic positive selection acting somewhere in the gene.

#### Site-specific pervasive selection: FUBAR

Fast Unconstrained Bayesian AppRoximation (FUBAR) was used to detect pervasive site-level selection acting consistently across lineages (Murrell et al., 2013). FUBAR estimates posterior probabilities for each codon site evolving under purifying selection (ω < 1) or diversifying selection (ω > 1) using a Bayesian framework. For each gene, FUBAR was run using default grid and MCMC settings. Codon sites with posterior probability ≥ 0.9 for ω > 1 were classified as being under pervasive positive selection, whereas sites with posterior probability ≥ 0.9 for ω < 1 were classified as being under pervasive purifying selection.

#### Amino-acid substitution bias: FADE

FUBAR Approach To Directional Evolution (FADE) tests whether substitutions along a specified lineage are directionally biased toward particular amino-acid residues (Pond et al., 2005). FADE requires a rooted tree, so trees were rooted at the split separating the monophyletic bat IAV clade from the remaining background taxa. FADE was run on protein alignments with the bat IAV clade designated as the test branch set. Directional amino-acid sites were evaluated using an empirical Bayes factor threshold of 100. Genes in which the bat-associated tips were not monophyletic under the required rooting scheme were not interpreted for FADE. FADE results were interpreted separately from codon-usage analyses because FADE tests directional amino-acid substitution bias, not synonymous codon bias.

#### Site-specific episodic selection: MEME

To identify episodic positive selection affecting individual codon sites on a subset of branches, we applied the Mixed Effects Model of Evolution (MEME) (Pond et al., 2005). Unlike FUBAR, MEME allows ω to vary across both sites and branches, enabling detection of transient selection events. MEME was run independently for each gene. Evidence for episodic positive selection at a codon site was determined using likelihood ratio tests, with sites considered significant at *p* < 0.05. MEME results were interpreted in conjunction with aBSREL to distinguish branch-specific versus site-specific episodic selection signals.

#### Branch-site selection: PAML

Branch-site models were implemented in PAML v4.10.10 to test for codons with elevated dN/dS on the bat-associated foreground lineage (Álvarez-Carretero et al., 2023; Yang, 1997). For each gene, a foreground branch corresponding to the bat IAV MRCA lineage was labeled per gene-specific tree. Model A, which allows a class of sites to have ω > 1 on the foreground branch, was compared with the corresponding null model in which ω was fixed at 1. Statistical significance was assessed using a likelihood-ratio test with a mixed χ^2^ distribution. Sites with elevated posterior support under Bayes Empirical Bayes analysis were extracted from significant PAML branch-site models as candidate foreground-associated sites (Yang et al., 2005).

### Codon-usage analysis

Codon-usage patterns were evaluated using the same reduced taxon set as selection analyses. Codon counts were extracted from the coding sequence alignments using coRdon (Elek, 2018). Relative synonymous codon usage (RSCU), GC content across the coding sequence (total GC content), GC content at the third codon positions (GC3), and the effective number of codons (ENC) were calculated for each sequence (Wright, 1990). Host-associated differences in multivariate RSCU profiles were tested by gene using PERMANOVA implemented with adonis2 in the R package vegan using Euclidean distances calculated from scaled RSCU values and 999 permutations (Anderson, 2001; McArdle & Anderson, 2001; Oksanen et al., 2026). Homogeneity of multivariate dispersion was evaluated using betadisper in the R package vegan (Anderson, 2006). Univariate differences in total GC content, GC3, ENC, and retained codon counts between bat and non-bat sequences were evaluated using Wilcoxon rank-sum tests, with Benjamini-Hochberg correction applied across tests (Wilcoxon, 1945; Benjamini & Hochberg, 1995). Individual codon-level RSCU differences were also evaluated using Wilcoxon rank-sum tests with Benjamini-Hochberg correction.

## Acknowledgements

The authors would like to thank Joe Manthey for helping with simplified host classifications. Lucas Ferreri, Siobain Duffy, Alvin Crespo-Bellido, and Madison Bullock for offering insights into pipeline development. The authors acknowledge the High Performance Computing Center (HPCC) at Texas Tech University for providing computational resources that have contributed to the research results reported within this paper. URL: http://www.hpcc.ttu.edu

## References

Álvarez-Carretero, S., Kapli, P., & Yang, Z. (2023). Beginner’s Guide on the Use of PAML to Detect Positive Selection. Molecular Biology and Evolution, 40(4), msad041. 10.1093/molbev/msad041

Anderson, M. J. (2001). A new method for non-parametric multivariate analysis of variance. Austral Ecology, 26(1), 32–46. 10.1111/j.1442-9993.2001.01070.pp.x

Anderson, M. J. (2006). Distance-based tests for homogeneity of multivariate dispersions. Biometrics, 62(1), 245–253. 10.1111/j.1541-0420.2005.00440.x

Ayres, D. L., Cummings, M. P., Baele, G., Darling, A. E., Lewis, P. O., Swofford, D. L., Huelsenbeck, J. P., Lemey, P., Rambaut, A., & Suchard, M. A. (2019). BEAGLE 3: Improved Performance, Scaling, and Usability for a High-Performance Computing Library for Statistical Phylogenetics. Systematic Biology, 68(6), 1052–1061. 10.1093/sysbio/syz020

Baele, G., Ji, X., Hassler, G. W., McCrone, J. T., Shao, Y., Zhang, Z., Holbrook, A. J., Lemey, P., Drummond, A. J., Rambaut, A., & Suchard, M. A. (2025). BEAST X for Bayesian phylogenetic, phylogeographic and phylodynamic inference. Nature Methods, 22(8), 1653–1656. 10.1038/s41592-025-02751-x

Bao, Y., Bolotov, P., Dernovoy, D., Kiryutin, B., Zaslavsky, L., Tatusova, T., Ostell, J., & Lipman, D. (2008). The Influenza Virus Resource at the National Center for Biotechnology Information. Journal of Virology, 82(2), 596–601. 10.1128/jvi.02005-07

Benjamini, Y., & Hochberg, Y. (1995). Controlling the False Discovery Rate: A Practical and Powerful Approach to Multiple Testing. Journal of the Royal Statistical Society. Series B (Methodological), 57(1), 289–300.

Brister, J. R., Ako-Adjei, D., Bao, Y., & Blinkova, O. (2015). NCBI Viral Genomes Resource. Nucleic Acids Research, 43(D1), D571–D577. 10.1093/nar/gku1207

Cardenas, M., Compton, S., Caceres, C. J., García-Sastre, A., Perez, D. R., & Rajao, D. S. (2026). MHC class II functions as a host-specific entry receptor for representative human and swine H3N2 influenza A viruses. Nature Communications, 17(1), 2560. 10.1038/s41467-026-69267-6

Chauché, C., Nogales, A., Zhu, H., Goldfarb, D., Ahmad Shanizza, A. I., Gu, Q., Parrish, C. R., Martínez-Sobrido, L., Marshall, J. F., & Murcia, P. R. (2018). Mammalian Adaptation of an Avian Influenza A Virus Involves Stepwise Changes in NS1. Journal of Virology, 92(5), 10.1128/jvi.01875-17. https://doi.org/10.1128/jvi.01875-17

Chothe, S. K., Bhushan, G., Nissly, R. H., Yeh, Y.-T., Brown, J., Turner, G., Fisher, J., Sewall, B. J., Reeder, D. M., Terrones, M., Jayarao, B. M., & Kuchipudi, S. V. (2017). Avian and human influenza virus compatible sialic acid receptors in little brown bats. Scientific Reports, 7(1), 660. 10.1038/s41598-017-00793-6

Ciminski, K., Pfaff, F., Beer, M., & Schwemmle, M. (2020). Bats reveal the true power of influenza A virus adaptability. PLoS Pathog, 16(4), e1008384. 10.1371/journal.ppat.1008384

Diskin, E. R., Friedman, K., Krauss, S., Nolting, J. M., Poulson, R. L., Slemons, R. D., Stallknecht, D. E., Webster, R. G., & Bowman, A. S. (2020). Subtype Diversity of Influenza A Virus in North American Waterfowl: A Multidecade Study. Journal of Virology, 94(11), 10.1128/jvi.02022-19. https://doi.org/10.1128/jvi.02022-19

Domingues, P., Eletto, D., Magnus, C., Turkington, H. L., Schmutz, S., Zagordi, O., Lenk, M., Beer, M., Stertz, S., & Hale, B. G. (2019). Profiling host ANP32A splicing landscapes to predict influenza A virus polymerase adaptation. Nature Communications, 10(1), 3396. 10.1038/s41467-019-11388-2

Echeverri-De la Hoz, D., Martínez-Bravo, C., Gastelbondo-Pastrana, B., Rivero, R., López, Y., Bertel, V., Alemán-Santos, M., Garay, E., Hoyos, R., Arrieta, G., Ramírez, J. D., & Mattar, S. (2025). Genomics of novel influenza A virus (H18N12) in bats, Caribe Colombia. Scientific Reports, 15(1), 6507. 10.1038/s41598-025-91026-8

Edwards, C. J., Suchard, M. A., Lemey, P., Welch, J. J., Barnes, I., Fulton, T. L., Barnett, R., O’Connell, T. C., Coxon, P., Monaghan, N., Valdiosera, C. E., Lorenzen, E. D., Willerslev, E., Baryshnikov, G. F., Rambaut, A., Thomas, M. G., Bradley, D. G., & Shapiro, B. (2011). Ancient hybridization and an Irish origin for the modern polar bear matriline. Current Biology, 21(15), 1251–1258. 10.1016/j.cub.2011.05.058

Elek, A. (2018, September 13). coRdon: An R package for codon usage analysis and prediction of gene expressivity. https://www.semanticscholar.org/paper/coRdon%3A-an-R-package-for-codon-usage-analysis-and-Elek/bb8e81a8b07eb6d67db6932e72544d81f2aea53d

Eriksson, P., Lindskog, C., Engholm, E., Blixt, O., Waldenstrom, J., Munster, V., Lundkvist, A., Olsen, B., Jourdain, E., & Ellstrom, P. (2018). Characterization of avian influenza virus attachment patterns to human and pig tissues. Sci Rep, 8(1), 12215. 10.1038/s41598-018-29578-1

Ferreri, L. M., Geiger, G., Seibert, B., Obadan, A., Rajao, D., Lowen, A. C., & Perez, D. R. (2022). Intra- and inter-host evolution of H9N2 influenza A virus in Japanese quail. Virus Evol, 8(1), veac001. 10.1093/ve/veac001

Ganti, K., Bagga, A., Carnaccini, S., Ferreri, L. M., Geiger, G., Joaquin Caceres, C., Seibert, B., Li, Y., Wang, L., Kwon, T., Li, Y., Morozov, I., Ma, W., Richt, J. A., Perez, D. R., Koelle, K., & Lowen, A. C. (2022). Influenza A virus reassortment in mammals gives rise to genetically distinct within-host subpopulations. Nature Communications, 13(1). 10.1038/s41467-022-34611-z

Glennon, N. B., Jabado, O., Lo, M. K., & Shaw, M. L. (2015). Transcriptome Profiling of the Virus-Induced Innate Immune Response in Pteropus vampyrus and Its Attenuation by Nipah Virus Interferon Antagonist Functions. Journal of Virology, 89(15), 7550–7566. 10.1128/jvi.00302-15

Hargadon, K. M., Zhou, H., Albrecht, R. A., Dodd, H. A., García-Sastre, A., & Braciale, T. J. (2011). Major Histocompatibility Complex Class II Expression and Hemagglutinin Subtype Influence the Infectivity of Type A Influenza Virus for Respiratory Dendritic Cells. Journal of Virology, 85(22), 11955–11963. 10.1128/jvi.05830-11

He, W., Li, G., Wang, R., Shi, W., Li, K., Wang, S., Lai, A., & Su, S. (2019). Host-range shift of H3N8 canine influenza virus: A phylodynamic analysis of its origin and adaptation from equine to canine host. Veterinary Research, 50(1), 87. 10.1186/s13567-019-0707-2

Hill, N. J., Bishop, M. A., Trovão, N. S., Ineson, K. M., Schaefer, A. L., Puryear, W. B., Zhou, K., Foss, A. D., Clark, D. E., MacKenzie, K. G., Jr, J. D. G., Borkenhagen, L. K., Hall, J. S., & Runstadler, J. A. (2022). Ecological divergence of wild birds drives avian influenza spillover and global spread. PLOS Pathogens, 18(5), e1010062. 10.1371/journal.ppat.1010062

Hockman, M. R., Phipps, K. L., Holmes, K. E., & Lowen, A. C. (2020). A method for the unbiased quantification of reassortment in segmented viruses. J Virol Methods, 280, 113878. 10.1016/j.jviromet.2020.113878

Holm, S. (1979). A Simple Sequentially Rejective Multiple Test Procedure. Scandinavian Journal of Statistics, 6(2), 65–70.

Irving, A. T., Ahn, M., Goh, G., Anderson, D. E., & Wang, L.-F. (2021). Lessons from the host defences of bats, a unique viral reservoir. Nature, 589(7842), 363–370. 10.1038/s41586-020-03128-0

Karakus, U., Mena, I., Kottur, J., Zahed, S. S. E., Seoane, R., Yildiz, S., Chen, L., Plancarte, M., Lindsay, L., Halpin, R., Stockwell, T. B., Wentworth, D. E., Boons, G.-J., Krammer, F., Stertz, S., Boyce, W., Vries, R. P. de, Aggarwal, A. K., & García-Sastre, A. (2024). H19 influenza A virus exhibits species-specific MHC class II receptor usage. Cell Host & Microbe, 32(7), 1089–1102.e10. 10.1016/j.chom.2024.05.018

Karakus, U., Pohl, M. O., & Stertz, S. (2020). Breaking the Convention: Sialoglycan Variants, Coreceptors, and Alternative Receptors for Influenza A Virus Entry. J Virol, 94(4). 10.1128/JVI.01357-19

Karakus, U., Sempere Borau, M., Martínez-Barragán, P., von Kempis, J., Yildiz, S., Arroyo-Fernández, L. M., Pohl, M. O., Steiger, J. A., Glas, I., Hunziker, A., García-Sastre, A., & Stertz, S. (2024). MHC class II proteins mediate sialic acid independent entry of human and avian H2N2 influenza A viruses. Nature Microbiology, 9(10), 2626–2641. 10.1038/s41564-024-01771-1

Karakus, U., Thamamongood, T., Ciminski, K., Ran, W., Günther, S. C., Pohl, M. O., Eletto, D., Jeney, C., Hoffmann, D., Reiche, S., Schinköthe, J., Ulrich, R., Wiener, J., Hayes, M. G. B., Chang, M. W., Hunziker, A., Yángüez, E., Aydillo, T., Krammer, F., … Stertz, S. (2019). MHC class II proteins mediate cross-species entry of bat influenza viruses. Nature, 567(7746), 109–112. 10.1038/s41586-019-0955-3

Kessler, S., Burke, B., Andrieux, G., Schinköthe, J., Hamberger, L., Kacza, J., Zhan, S., Reasoner, C., Dutt, T. S., Kaukab Osman, M., Henao-Tamayo, M., Staniek, J., Villena Ossa, J. F., Frank, D. T., Ma, W., Ulrich, R., Cathomen, T., Boerries, M., Rizzi, M., … Ciminski, K. (2024). Deciphering bat influenza H18N11 infection dynamics in male Jamaican fruit bats on a single-cell level. Nature Communications, 15(1), 4500. 10.1038/s41467-024-48934-6

Layan, M., Müller, N. F., Dellicour, S., De Maio, N., Bourhy, H., Cauchemez, S., & Baele, G. (2023). Impact and mitigation of sampling bias to determine viral spread: Evaluating discrete phylogeography through CTMC modeling and structured coalescent model approximations. Virus Evolution, 9(1), vead010. 10.1093/ve/vead010

Lemey, P., Rambaut, A., Drummond, A. J., & Suchard, M. A. (2009). Bayesian Phylogeography Finds Its Roots. PLOS Computational Biology, 5(9), e1000520. 10.1371/journal.pcbi.1000520

Li, Q., Sun, X., Li, Z., Liu, Y., Vavricka, C. J., Qi, J., & Gao, G. F. (2012). Structural and functional characterization of neuraminidase-like molecule N10 derived from bat influenza A virus. Proceedings of the National Academy of Sciences, 109(46), 18897–18902. 10.1073/pnas.1211037109

Maio, N. D., Wu, C.-H., O’Reilly, K. M., & Wilson, D. (2015). New Routes to Phylogeography: A Bayesian Structured Coalescent Approximation. PLOS Genetics, 11(8), e1005421. 10.1371/journal.pgen.1005421

Markin, A., Wagle, S., Grover, S., Vincent Baker, A. L., Eulenstein, O., & Anderson, T. K. (2023). PARNAS: Objectively Selecting the Most Representative Taxa on a Phylogeny. Systematic Biology, 72(5), 1052–1063. 10.1093/sysbio/syad028

McArdle, B. H., & Anderson, M. J. (2001). Fitting Multivariate Models to Community Data: A Comment on Distance-Based Redundancy Analysis. Ecology, 82(1), 290–297. 10.1890/0012-9658(2001)082%5B0290:FMMTCD%5D2.0.CO;2

Mikula, P., Morelli, F., Lučan, R. K., Jones, D. N., & Tryjanowski, P. (2016). Bats as prey of diurnal birds: A global perspective. Mammal Review, 46(3), 160–174. 10.1111/mam.12060

Minh, B. Q., Schmidt, H. A., Chernomor, O., Schrempf, D., Woodhams, M. D., von Haeseler, A., & Lanfear, R. (2020). IQ-TREE 2: New Models and Efficient Methods for Phylogenetic Inference in the Genomic Era. Molecular Biology and Evolution, 37(5), 1530–1534. 10.1093/molbev/msaa015

Murrell, B., Moola, S., Mabona, A., Weighill, T., Sheward, D., Kosakovsky Pond, S. L., & Scheffler, K. (2013). FUBAR: A fast, unconstrained bayesian approximation for inferring selection. Molecular Biology and Evolution, 30(5), 1196–1205. 10.1093/molbev/mst030

Murrell, B., Weaver, S., Smith, M. D., Wertheim, J. O., Murrell, S., Aylward, A., Eren, K., Pollner, T., Martin, D. P., Smith, D. M., Scheffler, K., & Kosakovsky Pond, S. L. (2015). Gene-Wide Identification of Episodic Selection. Molecular Biology and Evolution, 32(5), 1365–1371. 10.1093/molbev/msv035

Nabhan, A. R., & Sarkar, I. N. (2012). The impact of taxon sampling on phylogenetic inference: A review of two decades of controversy. Briefings in Bioinformatics, 13(1), 122–134. 10.1093/bib/bbr014

Oksanen, J., Simpson, G. L., Blanchet, F. G., Kindt, R., Legendre, P., Minchin, P. R., O’Hara, R. B., Solymos, P., Stevens, M. H. H., Szoecs, E., Wagner, H., Barbour, M., Bedward, M., Bengtsson, H., Bolker, B., Borcard, D., Borman, T., Carvalho, G., Chirico, M., … Weedon, J. (2026). vegan: Community Ecology Package (Version 2.7-5) [Computer software]. https://cran.r-project.org/web/packages/vegan/index.html

Parrish, C. R., Murcia, P. R., & Holmes, E. C. (2015). Influenza virus reservoirs and intermediate hosts: Dogs, horses, and new possibilities for influenza virus exposure of humans. J Virol, 89(6), 2990–2994. 10.1128/JVI.03146-14

Pond, S. L. K., Frost, S. D. W., & Muse, S. V. (2005). HyPhy: Hypothesis testing using phylogenies. Bioinformatics, 21(5), 676–679. 10.1093/bioinformatics/bti079

Price, M. N., Dehal, P. S., & Arkin, A. P. (2009). FastTree: Computing Large Minimum Evolution Trees with Profiles instead of a Distance Matrix. Molecular Biology and Evolution, 26(7), 1641–1650. 10.1093/molbev/msp077

Rambaut, A., Lam, T. T., Max Carvalho, L., & Pybus, O. G. (2016). Exploring the temporal structure of heterochronous sequences using TempEst (formerly Path-O-Gen). Virus Evolution, 2(1), vew007. 10.1093/ve/vew007

Ranwez, V., Douzery, E. J. P., Cambon, C., Chantret, N., & Delsuc, F. (2018). MACSE v2: Toolkit for the Alignment of Coding Sequences Accounting for Frameshifts and Stop Codons. Molecular Biology and Evolution, 35(10), 2582–2584. 10.1093/molbev/msy159

Rashid, F., Xie, Z., Li, M., Xie, Z., Luo, S., & Xie, L. (2023). Roles and functions of IAV proteins in host immune evasion. Frontiers in Immunology, 14, 1323560. 10.3389/fimmu.2023.1323560

Rognes, T., Flouri, T., Nichols, B., Quince, C., & Mahé, F. (2016). VSEARCH: A versatile open source tool for metagenomics. PeerJ, 4, e2584. 10.7717/peerj.2584

Smith, M. D., Wertheim, J. O., Weaver, S., Murrell, B., Scheffler, K., & Kosakovsky Pond, S. L. (2015). Less Is More: An Adaptive Branch-Site Random Effects Model for Efficient Detection of Episodic Diversifying Selection. Molecular Biology and Evolution, 32(5), 1342–1353. 10.1093/molbev/msv022

Sun, X., Shi, Y., Lu, X., He, J., Gao, F., Yan, J., Qi, J., & Gao, G. F. (2013). Bat-Derived Influenza Hemagglutinin H17 Does Not Bind Canonical Avian or Human Receptors and Most Likely Uses a Unique Entry Mechanism. Cell Reports, 3(3), 769–778. 10.1016/j.celrep.2013.01.025

Tavaré, S. (1986). Some probabilistic and statistical problems in the analysis of DNA sequences. https://www.semanticscholar.org/paper/Some-probabilistic-and-statistical-problems-in-the-Tavar%C3%A9/55e3359cd05b1903ffd8f633eb5aa6156791b364

Tisoncik, J. R., Billharz, R., Burmakina, S., Belisle, S. E., Proll, S. C., Korth, M. J., García-Sastre, A., & Katze, M. G. (2011). The NS1 protein of influenza A virus suppresses interferon-regulated activation of antigen-presentation and immune-proteasome pathways. The Journal of General Virology, 92(Pt 9), 2093–2104. 10.1099/vir.0.032060-0

Tong, S., Li, Y., Rivailler, P., Conrardy, C., Castillo, D. A. A., Chen, L.-M., Recuenco, S., Ellison, J. A., Davis, C. T., York, I. A., Turmelle, A. S., Moran, D., Rogers, S., Shi, M., Tao, Y., Weil, M. R., Tang, K., Rowe, L. A., Sammons, S., … Donis, R. O. (2012). A distinct lineage of influenza A virus from bats. Proceedings of the National Academy of Sciences, 109(11), 4269–4274. 10.1073/pnas.1116200109

Turkington, H. L., Juozapaitis, M., Tsolakos, N., Corrales-Aguilar, E., Schwemmle, M., & Hale, B. G. (2018). Unexpected Functional Divergence of Bat Influenza Virus NS1 Proteins. Journal of Virology, 92(5), 10.1128/jvi.02097-17. https://doi.org/10.1128/jvi.02097-17

Webby, R. J., Swenson, S. L., Krauss, S. L., Gerrish, P. J., Goyal, S. M., & Webster, R. G. (2000). Evolution of Swine H3N2 Influenza Viruses in the United States. Journal of Virology, 74(18), 8243–8251. 10.1128/jvi.74.18.8243-8251.2000

Wertheim, J. O., Murrell, B., Smith, M. D., Kosakovsky Pond, S. L., & Scheffler, K. (2015). RELAX: Detecting Relaxed Selection in a Phylogenetic Framework. Molecular Biology and Evolution, 32(3), 820–832. 10.1093/molbev/msu400

Wilcoxon, F. (1945). Individual Comparisons by Ranking Methods. Biometrics Bulletin, 1(6), 80–83. 10.2307/3001968

Wolf, Y. I., Viboud, C., Holmes, E. C., Koonin, E. V., & Lipman, D. J. (2006). Long intervals of stasis punctuated by bursts of positive selection in the seasonal evolution of influenza A virus. Biology Direct, 1(1), 34. 10.1186/1745-6150-1-34

Wright, F. (1990). The “effective number of codons” used in a gene. Gene, 87(1), 23–29. 10.1016/0378-1119(90)90491-9

Wu, Y., Wu, Y., Tefsen, B., Shi, Y., & Gao, G. F. (2014). Bat-derived influenza-like viruses H17N10 and H18N11. Trends Microbiol, 22(4), 183–191. 10.1016/j.tim.2014.01.010

Xie, J., Li, Y., Shen, X., Goh, G., Zhu, Y., Cui, J., Wang, L.-F., Shi, Z.-L., & Zhou, P. (2018). Dampened STING-Dependent Interferon Activation in Bats. Cell Host & Microbe, 23(3), 297–301.e4. 10.1016/j.chom.2018.01.006

Yang, Z. (1994). Maximum likelihood phylogenetic estimation from DNA sequences with variable rates over sites: Approximate methods. Journal of Molecular Evolution, 39(3), 306–314. 10.1007/BF00160154

Yang, Z. (1997). PAML: A program package for phylogenetic analysis by maximum likelihood. Computer Applications in the Biosciences: CABIOS, 13(5), 555–556. 10.1093/bioinformatics/13.5.555

Yang, Z., & Bielawski, J. P. (2000). Statistical methods for detecting molecular adaptation. Trends in Ecology & Evolution, 15(12), 496–503. 10.1016/S0169-5347(00)01994-7

Yang, Z., Wong, W. S. W., & Nielsen, R. (2005). Bayes Empirical Bayes Inference of Amino Acid Sites Under Positive Selection. Molecular Biology and Evolution, 22(4), 1107–1118. 10.1093/molbev/msi097

Zwickl, D. J., & Hillis, D. M. (2002). Increased taxon sampling greatly reduces phylogenetic error. Systematic Biology, 51(4), 588–598. 10.1080/10635150290102339

