## Supplemental figures for "Evolutionary reconstruction of bat influenza A viruses reveals segment-specific histories and host-associated selection"

### Supplemental information

***Supplemental Table 1: LSD2 tMRCA estimates for bat-associated influenza lineages in the expanded dataset and 75-taxon subset.*** *Dates are reported as estimated calendar years for the bat-associated MRCA. Confidence intervals are node-specific LSD2 date intervals. UFBoot values indicate ultrafast bootstrap support for the corresponding MRCA node. NS1 and M2 subset estimates were classified as unstable because reduced-tree dating produced biologically implausible dates and extreme confidence intervals.*

| Gene | Expanded dataset | | | 75-taxon subset | | |
|---|---|---|---|---|---|---|
| | tMRCA | LSD2 CI | UFBoot | tMRCA | LSD2 CI | UFBoot |
| **Polymerase PB2** | 1932 | 1920–1942 | 100 | 1899 | 1880–1914 | 100 |
| **Polymerase PB1** | 1871 | 1837–1901 | 100 | 1678 | 1543–1766 | 100 |
| **Polymerase PA** | 1795 | 1736–1846 | 100 | 1661 | 1535–1754 | 99 |
| **Hemagglutinin (HA)** | 1830 | 1818–1840 | 100 | 1730 | 1613–1805 | 100 |
| **Nucleocapsid (NP)** | 1840 | 1797–1874 | 98 | 1837 | 1790–1874 | 74 |
| **Neuraminidase (NA)** | 1532 | 1428–1604 | 100 | 1440 | 987–1632 | 100 |
| **Matrix protein 1 (M1)** | 1880 | 1859–1893 | 97 | 1898 | 1867–1917 | 97 |
| **Non-structural protein 1 (NS1)** | 1861 | 1824–1898 | 100 | Unstable | Unstable | 100 |
| **Matrix protein 2 (M2)** | 1916 | 1876–1938 | 100 | Unstable | Unstable | 100 |
| **Nuclear export protein (NEP)** | 1924 | 1766–1972 | 100 | 1962 | 1925–1987 | 100 |

***Supplemental Table 2: PAML branch-site selection results for bat-associated influenza gene segments.*** *Likelihood-ratio test statistics and p-values compare Model A with the null model, while Bayes Empirical Bayes (BEB) posterior probabilities identify candidate codon sites associated with positive selection on the bat-associated foreground lineage.*

| Gene | Branch-site LRT statistic (2ΔlnL) | Branch-site LRT p-value | BEB posterior probability | Model A log-likelihood | Null model log-likelihood | BEB candidate site |
|---|---|---|---|---|---|---|
| PB1 (polymerase PB1) | 53.336502 | 1.41E-13 | 0.985 | -48527.12764 | -48553.79589 | 120N |
| PB1 (polymerase PB1) | 53.336502 | 1.41E-13 | 0.966 | -48527.12764 | -48553.79589 | 202E |
| PB1 (polymerase PB1) | 53.336502 | 1.41E-13 | 0.998 | -48527.12764 | -48553.79589 | 208E |
| PB1 (polymerase PB1) | 53.336502 | 1.41E-13 | 0.992 | -48527.12764 | -48553.79589 | 370S |
| PB1 (polymerase PB1) | 53.336502 | 1.41E-13 | 0.965 | -48527.12764 | -48553.79589 | 406A |
| PB1 (polymerase PB1) | 53.336502 | 1.41E-13 | 0.97 | -48527.12764 | -48553.79589 | 458S |
| PB1 (polymerase PB1) | 53.336502 | 1.41E-13 | 0.999 | -48527.12764 | -48553.79589 | 470Y |
| PB1 (polymerase PB1) | 53.336502 | 1.41E-13 | 0.994 | -48527.12764 | -48553.79589 | 494N |
| PB1 (polymerase PB1) | 53.336502 | 1.41E-13 | 0.998 | -48527.12764 | -48553.79589 | 519S |

| | | | | | | |
|---|---|---|---|---|---|---|
| PB1 (polymerase PB1) | 53.336502 | 1.41E-13 | 0.971 | -48527.12764 | -48553.79589 | 624E |
| PB1 (polymerase PB1) | 53.336502 | 1.41E-13 | 0.95 | -48527.12764 | -48553.79589 | 676S |
| PB1 (polymerase PB1) | 53.336502 | 1.41E-13 | 0.989 | -48527.12764 | -48553.79589 | 685N |
| PB1 (polymerase PB1) | 53.336502 | 1.41E-13 | 0.99 | -48527.12764 | -48553.79589 | 698S |
| PB1 (polymerase PB1) | 53.336502 | 1.41E-13 | 0.982 | -48527.12764 | -48553.79589 | 741N |
| PB1 (polymerase PB1) | 53.336502 | 1.41E-13 | 0.96 | -48527.12764 | -48553.79589 | 806S |
| PA (polymerase PA) | 60.46694 | 3.74E-15 | 0.982 | -45962.37286 | -45992.60633 | 255S |
| PA (polymerase PA) | 60.46694 | 3.74E-15 | 0.962 | -45962.37286 | -45992.60633 | 265D |
| PA (polymerase PA) | 60.46694 | 3.74E-15 | 0.964 | -45962.37286 | -45992.60633 | 290P |
| PA (polymerase PA) | 60.46694 | 3.74E-15 | 0.952 | -45962.37286 | -45992.60633 | 296P |
| PA (polymerase PA) | 60.46694 | 3.74E-15 | 0.97 | -45962.37286 | -45992.60633 | 309P |
| PA (polymerase PA) | 60.46694 | 3.74E-15 | 0.958 | -45962.37286 | -45992.60633 | 672S |
| HA (hemagglutinin) | 9.276692 | 1.16E-03 | 0.961 | -70720.01646 | -70724.6548 | 217N |
| HA (hemagglutinin) | 9.276692 | 1.16E-03 | 0.951 | -70720.01646 | -70724.6548 | 284V |
| NP (nucleocapsid) | 46.509396 | 4.56E-12 | 1 | -32953.38883 | -32976.64353 | 64S |
| NP (nucleocapsid) | 46.509396 | 4.56E-12 | 0.996 | -32953.38883 | -32976.64353 | 75K |
| NP (nucleocapsid) | 46.509396 | 4.56E-12 | 0.998 | -32953.38883 | -32976.64353 | 90V |
| NP (nucleocapsid) | 46.509396 | 4.56E-12 | 0.985 | -32953.38883 | -32976.64353 | 106N |
| NP (nucleocapsid) | 46.509396 | 4.56E-12 | 0.998 | -32953.38883 | -32976.64353 | 112N |
| NP (nucleocapsid) | 46.509396 | 4.56E-12 | 1 | -32953.38883 | -32976.64353 | 179F |
| NP (nucleocapsid) | 46.509396 | 4.56E-12 | 0.986 | -32953.38883 | -32976.64353 | 183E |
| NP (nucleocapsid) | 46.509396 | 4.56E-12 | 0.993 | -32953.38883 | -32976.64353 | 186T |
| NP (nucleocapsid) | 46.509396 | 4.56E-12 | 0.996 | -32953.38883 | -32976.64353 | 189K |
| NP (nucleocapsid) | 46.509396 | 4.56E-12 | 0.999 | -32953.38883 | -32976.64353 | 197G |
| NP (nucleocapsid) | 46.509396 | 4.56E-12 | 0.998 | -32953.38883 | -32976.64353 | 198F |
| NP (nucleocapsid) | 46.509396 | 4.56E-12 | 1 | -32953.38883 | -32976.64353 | 357I |
| NA (neuraminidase) | 13.848784 | 9.91E-05 | 0.999 | -56444.64305 | -56451.56744 | 20N |
| NA (neuraminidase) | 13.848784 | 9.91E-05 | 0.97 | -56444.64305 | -56451.56744 | 21P |

| | | | | | | |
|---|---|---|---|---|---|---|
| NA (neuraminidase) | 13.848784 | 9.91E-05 | 0.958 | -56444.64305 | -56451.56744 | 25Q |
| NA (neuraminidase) | 13.848784 | 9.91E-05 | 0.953 | -56444.64305 | -56451.56744 | 26K |
| NA (neuraminidase) | 13.848784 | 9.91E-05 | 0.999 | -56444.64305 | -56451.56744 | 31F |
| NA (neuraminidase) | 13.848784 | 9.91E-05 | 0.996 | -56444.64305 | -56451.56744 | 32A |
| NA (neuraminidase) | 13.848784 | 9.91E-05 | 0.99 | -56444.64305 | -56451.56744 | 54L |
| NA (neuraminidase) | 13.848784 | 9.91E-05 | 0.994 | -56444.64305 | -56451.56744 | 58S |
| NA (neuraminidase) | 13.848784 | 9.91E-05 | 0.982 | -56444.64305 | -56451.56744 | 59N |
| NA (neuraminidase) | 13.848784 | 9.91E-05 | 0.952 | -56444.64305 | -56451.56744 | 60V |
| NA (neuraminidase) | 13.848784 | 9.91E-05 | 0.98 | -56444.64305 | -56451.56744 | 124S |
| NA (neuraminidase) | 13.848784 | 9.91E-05 | 0.991 | -56444.64305 | -56451.56744 | 203I |
| NA (neuraminidase) | 13.848784 | 9.91E-05 | 0.997 | -56444.64305 | -56451.56744 | 218S |
| NA (neuraminidase) | 13.848784 | 9.91E-05 | 0.982 | -56444.64305 | -56451.56744 | 257T |
| NA (neuraminidase) | 13.848784 | 9.91E-05 | 0.997 | -56444.64305 | -56451.56744 | 258T |
| NA (neuraminidase) | 13.848784 | 9.91E-05 | 0.989 | -56444.64305 | -56451.56744 | 259I |
| NA (neuraminidase) | 13.848784 | 9.91E-05 | 0.971 | -56444.64305 | -56451.56744 | 264H |
| NA (neuraminidase) | 13.848784 | 9.91E-05 | 0.977 | -56444.64305 | -56451.56744 | 267G |
| NA (neuraminidase) | 13.848784 | 9.91E-05 | 1 | -56444.64305 | -56451.56744 | 279I |
| NA (neuraminidase) | 13.848784 | 9.91E-05 | 0.998 | -56444.64305 | -56451.56744 | 311S |
| NA (neuraminidase) | 13.848784 | 9.91E-05 | 0.975 | -56444.64305 | -56451.56744 | 361K |
| NA (neuraminidase) | 13.848784 | 9.91E-05 | 0.983 | -56444.64305 | -56451.56744 | 364W |
| NA (neuraminidase) | 13.848784 | 9.91E-05 | 0.997 | -56444.64305 | -56451.56744 | 368I |
| NA (neuraminidase) | 13.848784 | 9.91E-05 | 0.993 | -56444.64305 | -56451.56744 | 425E |
| NA (neuraminidase) | 13.848784 | 9.91E-05 | 0.998 | -56444.64305 | -56451.56744 | 466E |
| NA (neuraminidase) | 13.848784 | 9.91E-05 | 0.96 | -56444.64305 | -56451.56744 | 483V |
| NA (neuraminidase) | 13.848784 | 9.91E-05 | 0.999 | -56444.64305 | -56451.56744 | 508N |
| NA (neuraminidase) | 13.848784 | 9.91E-05 | 0.995 | -56444.64305 | -56451.56744 | 539Y |
| NA (neuraminidase) | 13.848784 | 9.91E-05 | 0.979 | -56444.64305 | -56451.56744 | 540V |
| NA (neuraminidase) | 13.848784 | 9.91E-05 | 0.998 | -56444.64305 | -56451.56744 | 551P |

| | | | | | | |
|---|---|---|---|---|---|---|
| NA (neuraminidase) | 13.848784 | 9.91E-05 | 0.99 | -56444.64305 | -56451.56744 | 565P |
| NA (neuraminidase) | 13.848784 | 9.91E-05 | 0.988 | -56444.64305 | -56451.56744 | 578P |
| NA (neuraminidase) | 13.848784 | 9.91E-05 | 0.968 | -56444.64305 | -56451.56744 | 587G |
| NA (neuraminidase) | 13.848784 | 9.91E-05 | 0.981 | -56444.64305 | -56451.56744 | 642K |
| NA (neuraminidase) | 13.848784 | 9.91E-05 | 0.991 | -56444.64305 | -56451.56744 | 651Q |
| NA (neuraminidase) | 13.848784 | 9.91E-05 | 0.999 | -56444.64305 | -56451.56744 | 662Y |
| NA (neuraminidase) | 13.848784 | 9.91E-05 | 0.993 | -56444.64305 | -56451.56744 | 667S |
| NA (neuraminidase) | 13.848784 | 9.91E-05 | 0.999 | -56444.64305 | -56451.56744 | 670I |
| NA (neuraminidase) | 13.848784 | 9.91E-05 | 0.978 | -56444.64305 | -56451.56744 | 690Y |
| NA (neuraminidase) | 13.848784 | 9.91E-05 | 0.991 | -56444.64305 | -56451.56744 | 725E |
| NA (neuraminidase) | 13.848784 | 9.91E-05 | 0.968 | -56444.64305 | -56451.56744 | 743L |
| NA (neuraminidase) | 13.848784 | 9.91E-05 | 0.998 | -56444.64305 | -56451.56744 | 754P |
| NA (neuraminidase) | 13.848784 | 9.91E-05 | 0.998 | -56444.64305 | -56451.56744 | 758G |
| NA (neuraminidase) | 13.848784 | 9.91E-05 | 0.998 | -56444.64305 | -56451.56744 | 763S |
| M1 (matrix protein 1) | 9.642898 | 9.50E-04 | 0.979 | -16633.85604 | -16638.67749 | 79L |
| M1 (matrix protein 1) | 9.642898 | 9.50E-04 | 0.952 | -16633.85604 | -16638.67749 | 96N |
| M1 (matrix protein 1) | 9.642898 | 9.50E-04 | 0.953 | -16633.85604 | -16638.67749 | 197L |
| M1 (matrix protein 1) | 9.642898 | 9.50E-04 | 0.972 | -16633.85604 | -16638.67749 | 199S |
| M1 (matrix protein 1) | 9.642898 | 9.50E-04 | 0.998 | -16633.85604 | -16638.67749 | 212S |
| M1 (matrix protein 1) | 9.642898 | 9.50E-04 | 0.993 | -16633.85604 | -16638.67749 | 243S |
| NS1 (nonstructural protein 1) | 23.95368 | 4.93E-07 | 0.97 | -18151.12376 | -18163.1006 | 120Y |
| NS1 (nonstructural protein 1) | 23.95368 | 4.93E-07 | 0.975 | -18151.12376 | -18163.1006 | 122G |
| NS1 (nonstructural protein 1) | 23.95368 | 4.93E-07 | 0.972 | -18151.12376 | -18163.1006 | 128G |
| NS1 (nonstructural protein 1) | 23.95368 | 4.93E-07 | 0.995 | -18151.12376 | -18163.1006 | 135S |
| NS1 (nonstructural protein 1) | 23.95368 | 4.93E-07 | 0.971 | -18151.12376 | -18163.1006 | 167Q |

| | | | | | | |
|---|---|---|---|---|---|---|
| NS1 (nonstructural protein 1) | 23.95368 | 4.93E-07 | 0.996 | -18151.12376 | -18163.1006 | 236N |
| NS1 (nonstructural protein 1) | 23.95368 | 4.93E-07 | 0.957 | -18151.12376 | -18163.1006 | 257Y |
| NS1 (nonstructural protein 1) | 23.95368 | 4.93E-07 | 0.959 | -18151.12376 | -18163.1006 | 303P |
| M2 (matrix protein 2) | 9.966566 | 7.97E-04 | 0.953 | -7326.145742 | -7331.129025 | 82L |
| M2 (matrix protein 2) | 9.966566 | 7.97E-04 | 0.993 | -7326.145742 | -7331.129025 | 163S |
| M2 (matrix protein 2) | 9.966566 | 7.97E-04 | 0.992 | -7326.145742 | -7331.129025 | 236D |
| NEP (nuclear export protein) | 9.151608 | 1.24E-03 | 0.996 | -6200.416558 | -6204.992362 | 183S |
| NEP (nuclear export protein) | 9.151608 | 1.24E-03 | 0.995 | -6200.416558 | -6204.992362 | 217Q |

***Supplemental Table 43: Gene-specific comparisons of codon-bias metrics between bat and non-bat influenza sequences.*** *GC3, total GC content, and effective number of codons were compared using Wilcoxon rank-sum tests. Mean and median values summarize the direction of each host-associated difference, while q-values indicate Benjamini–Hochberg-corrected significance. Significant differences were primarily observed for GC-related metrics, suggesting that host-associated codon usage variation is partly associated with nucleotide composition at synonymous sites.*

| gene | metric | n groups | mean Bat | median Bat | mean Non bat | median Non bat | statistic | p value | q value BH within metric | q value BH all metric tests |
|---|---|---|---|---|---|---|---|---|---|---|
| PB2 | GC3 | 2 | 0.35 | 0.31 | 0.44 | 0.45 | 35.00 | 0.05 | 0.11 | 0.12 |
| PB2 | GC_total | 2 | 0.39 | 0.37 | 0.44 | 0.44 | 13.00 | 0.01 | 0.03 | **0.04** |
| PB2 | ENC_cordon | 2 | 47.46 | 45.75 | 50.98 | 51.25 | 43.00 | 0.08 | 0.27 | 0.15 |
| PB1 | GC3 | 2 | 0.35 | 0.30 | 0.44 | 0.45 | 35.00 | 0.05 | 0.11 | 0.12 |
| PB1 | GC_total | 2 | 0.38 | 0.36 | 0.43 | 0.43 | 39.00 | 0.06 | 0.11 | 0.13 |
| PB1 | ENC_cordon | 2 | 46.30 | 44.06 | 51.13 | 51.00 | 37.00 | 0.06 | 0.27 | 0.12 |
| PA | GC3 | 2 | 0.33 | 0.29 | 0.45 | 0.46 | 14.00 | 0.01 | 0.06 | **0.04** |
| PA | GC_total | 2 | 0.37 | 0.35 | 0.43 | 0.43 | 9.00 | 0.01 | 0.03 | **0.04** |
| PA | ENC_cordon | 2 | 45.26 | 42.84 | 51.96 | 52.53 | 22.00 | 0.02 | 0.21 | 0.06 |
| HA | GC3 | 2 | 0.40 | 0.40 | 0.42 | 0.42 | 60.00 | 0.68 | 0.85 | 0.80 |
| HA | GC_total | 2 | 0.40 | 0.40 | 0.42 | 0.42 | 34.50 | 0.21 | 0.26 | 0.33 |
| HA | ENC_cordon | 2 | 48.74 | 48.74 | 50.92 | 51.48 | 47.00 | 0.40 | 0.73 | 0.60 |
| NP | GC3 | 2 | 0.44 | 0.44 | 0.45 | 0.46 | 68.00 | 0.88 | 0.98 | 0.98 |
| NP | GC_total | 2 | 0.45 | 0.45 | 0.47 | 0.47 | 28.00 | 0.14 | 0.20 | 0.25 |
| NP | ENC_cordon | 2 | 50.90 | 50.90 | 49.92 | 50.01 | 97.00 | 0.44 | 0.73 | 0.63 |
| NA | GC3 | 2 | 0.41 | 0.42 | 0.41 | 0.41 | 143.00 | 0.99 | 0.99 | 1.00 |
| NA | GC_total | 2 | 0.43 | 0.43 | 0.43 | 0.43 | 119.00 | 0.60 | 0.64 | 0.77 |
| NA | ENC_cordon | 2 | 51.94 | 51.87 | 51.83 | 52.13 | 129.00 | 0.77 | 1.00 | 0.88 |
| M1 | GC3 | 2 | 0.46 | 0.46 | 0.50 | 0.51 | 37.00 | 0.06 | 0.11 | 0.12 |
| M1 | GC_total | 2 | 0.46 | 0.45 | 0.49 | 0.49 | 13.00 | 0.01 | 0.03 | **0.04** |
| M1 | ENC_cordon | 2 | 54.63 | 54.45 | 53.92 | 54.74 | 105.00 | 0.95 | 1.00 | 1.00 |
| M2 | GC3 | 2 | 0.39 | 0.38 | 0.46 | 0.45 | 13.00 | 0.01 | 0.06 | **0.04** |
| M2 | GC_total | 2 | 0.39 | 0.38 | 0.45 | 0.45 | 2.00 | 0.00 | 0.03 | **0.03** |
| M2 | ENC_cordon | 2 | 49.20 | 51.78 | 51.22 | 50.52 | 108.00 | 1.00 | 1.00 | 1.00 |
| NS1 | GC3 | 2 | 0.38 | 0.37 | 0.42 | 0.42 | 58.00 | 0.18 | 0.30 | 0.29 |
| NS1 | GC_total | 2 | 0.43 | 0.42 | 0.45 | 0.46 | 22.00 | 0.02 | 0.04 | 0.06 |
| NS1 | ENC_cordon | 2 | 48.37 | 46.99 | 51.72 | 50.54 | 53.00 | 0.14 | 0.35 | 0.25 |
| NS2/NEP | GC3 | 2 | 0.48 | 0.48 | 0.48 | 0.47 | 70.00 | 0.67 | 0.85 | 0.80 |
| NS2/NEP | GC_total | 2 | 0.40 | 0.40 | 0.41 | 0.41 | 47.00 | 0.64 | 0.64 | 0.80 |
| NS2/NEP | ENC_cordon | 2 | 48.66 | 48.66 | 48.89 | 48.17 | 59.00 | 1.00 | 1.00 | 1.00 |

***Supplemental Table 4: NCBI Virus accession list for each gene with encoded host state and date used for divergence estimates, selection signal, codon usage, and ancestral host reconstruction.***

| Accession | Gene | Segment | Encoded host state | Date |
|---|---|---|---|---|
| PX457967.1 | PB2 | segment_1 | Avian | 2022 |
| MW871633.1 | PB2 | segment_1 | Avian | 2018 |
| DQ208309.1 | PB2 | segment_1 | Avian | 1918 |
| CY005822.1 | PB2 | segment_1 | Avian | 1956 |
| MN530188.1 | PB2 | segment_1 | Avian | 2012 |
| CY003862.1 | PB2 | segment_1 | Avian | 1977 |
| CY014686.1 | PB2 | segment_1 | Avian | 1956 |
| EU871835.1 | PB2 | segment_1 | Avian | 1987 |
| CY179706.1 | PB2 | segment_1 | Avian | 1975 |
| GU052882.1 | PB2 | segment_1 | Avian | 1963 |
| AB274963.1 | PB2 | segment_1 | Human | 1980 |
| PV946091.1 | PB2 | segment_1 | Avian | 2024 |
| GU052875.1 | PB2 | segment_1 | Avian | 2002 |
| CY005819.1 | PB2 | segment_1 | Avian | 1963 |
| MW094698.1 | PB2 | segment_1 | Avian | 2017 |

| | | | | |
|---|---|---|---|---|
| CY005624.1 | PB2 | segment_1 | Avian | 1978 |
| KF619244.1 | PB2 | segment_1 | Avian | 1925 |
| PV306261.1 | PB2 | segment_1 | Avian | 2022 |
| PV370541.1 | PB2 | segment_1 | Avian | 2022 |
| OQ861242.1 | PB2 | segment_1 | Avian | 2022 |
| MH500872.1 | PB2 | segment_1 | Avian | 2001 |
| LC041310.1 | PB2 | segment_1 | Avian | 2014 |
| GU051972.1 | PB2 | segment_1 | Avian | 1966 |
| CY014998.1 | PB2 | segment_1 | Avian | 1927 |
| CY014693.1 | PB2 | segment_1 | Avian | 1974 |
| CY130037.1 | PB2 | segment_1 | Avian | 1965 |
| ON326607.1 | PB2 | segment_1 | Avian | 2020 |
| PQ579025.1 | PB2 | segment_1 | Avian | 2021 |
| MK996233.1 | PB2 | segment_1 | Avian | 2019 |
| KP185514.1 | PB2 | segment_1 | Avian | 2013 |
| CY015053.1 | PB2 | segment_1 | Avian | 1934 |
| CY067691.1 | PB2 | segment_1 | Avian | 2007 |
| MH134703.1 | PB2 | segment_1 | Avian | 2016 |
| MH498899.1 | PB2 | segment_1 | Avian | 2017 |
| KX185897.1 | PB2 | segment_1 | Avian | 2015 |
| OQ861233.1 | PB2 | segment_1 | Avian | 2022 |
| JX175250.1 | PB2 | segment_1 | Avian | 2011 |
| OP476926.1 | PB2 | segment_1 | Avian | 2020 |
| CY028843.1 | PB2 | segment_1 | Non-human mammal | 1963 |
| LC364318.1 | PB2 | segment_1 | Avian | 2017 |
| MF694187.1 | PB2 | segment_1 | Avian | 2016 |
| MW090832.1 | PB2 | segment_1 | Avian | 2016 |
| PV739618.1 | PB2 | segment_1 | Avian | 2025 |
| JX494721.1 | PB2 | segment_1 | Non-human mammal | 2010 |
| OP535288.1 | PB2 | segment_1 | Avian | 2022 |
| PV235955.1 | PB2 | segment_1 | Avian | 2024 |
| MK361511.1 | PB2 | segment_1 | Non-human mammal | 1987 |
| PP345568.1 | PB2 | segment_1 | Avian | 2019 |
| MT666559.1 | PB2 | segment_1 | Non-human mammal | 2018 |
| PQ191259.1 | PB2 | segment_1 | Avian | 2023 |
| PP653396.1 | PB2 | segment_1 | Human | 2024 |
| MW295727.1 | PB2 | segment_1 | Non-human mammal | 2013 |
| CY039398.1 | PB2 | segment_1 | Non-human mammal | 1966 |
| MF116344.1 | PB2 | segment_1 | Avian | 2012 |
| PV770031.1 | PB2 | segment_1 | Human | 2024 |
| FR687270.1 | PB2 | segment_1 | Avian | 2010 |

| | | | | |
|---|---|---|---|---|
| PP929863.1 | PB2 | segment_1 | Avian | 2024 |
| MN882398.1 | PB2 | segment_1 | Avian | 2017 |
| PP664071.1 | PB2 | segment_1 | Environment | 2022 |
| PQ356534.1 | PB2 | segment_1 | Human | 2024 |
| AB600245.1 | PB2 | segment_1 | Non-human mammal | 2004 |
| FJ839891.1 | PB2 | segment_1 | Non-human mammal | 2002 |
| MG962429.1 | PB2 | segment_1 | Non-human mammal | 2012 |
| OR762271.1 | PB2 | segment_1 | Non-human mammal | 2021 |
| JN043464.1 | PB2 | segment_1 | Non-human mammal | 2009 |
| PV388416.1 | PB2 | segment_1 | Avian | 2024 |
| KF881680.1 | PB2 | segment_1 | Avian | 2013 |
| AB573797.1 | PB2 | segment_1 | Non-human mammal | 2005 |
| MH376909.1 | PB2 | segment_1 | Bat | 2017 |
| OQ674405.1 | PB2 | segment_1 | Avian | 2022 |
| CY125942.1 | PB2 | segment_1 | Bat | 2010 |
| MH682200.1 | PB2 | segment_1 | Bat | 2012 |
| CY103873.1 | PB2 | segment_1 | Bat | 2009 |
| PV233913.1 | PB2 | segment_1 | Human | 2023 |
| DQ454102.2 | PB2 | segment_1 | Avian | 2004 |
| PX461673.1 | PB1 | segment_2 | Avian | 2022 |
| PX444270.1 | PB1 | segment_2 | Non-human mammal | 2021 |
| PQ115506.1 | PB1 | segment_2 | Non-human mammal | 2020 |
| PV412341.1 | PB1 | segment_2 | Avian | 2021 |
| CY015045.1 | PB1 | segment_2 | Avian | 1979 |
| GU361340.1 | PB1 | segment_2 | Avian | 2007 |
| PQ675394.1 | PB1 | segment_2 | Avian | 2024 |
| MH592195.1 | PB1 | segment_2 | Avian | 2012 |
| CY005799.1 | PB1 | segment_2 | Avian | 1965 |
| PP333639.1 | PB1 | segment_2 | Avian | 2023 |
| CY014669.1 | PB1 | segment_2 | Avian | 1966 |
| CY014997.1 | PB1 | segment_2 | Avian | 1927 |
| CY004590.1 | PB1 | segment_2 | Avian | 1977 |
| CY179865.1 | PB1 | segment_2 | Avian | 1975 |
| CY004120.1 | PB1 | segment_2 | Avian | 1979 |
| CY015087.1 | PB1 | segment_2 | Avian | 1959 |
| KF619245.1 | PB1 | segment_2 | Avian | 1925 |
| KF435047.1 | PB1 | segment_2 | Avian | 1953 |
| OQ862272.1 | PB1 | segment_2 | Avian | 2022 |
| KF193392.1 | PB1 | segment_2 | Avian | 2012 |
| MW265245.1 | PB1 | segment_2 | Avian | 2018 |
| CY005821.1 | PB1 | segment_2 | Avian | 1956 |

| | | | | |
|---|---|---|---|---|
| CY015052.1 | PB1 | segment_2 | Avian | 1934 |
| MW205010.1 | PB1 | segment_2 | Avian | 2017 |
| MK806578.1 | PB1 | segment_2 | Non-human mammal | 2019 |
| KY925620.1 | PB1 | segment_2 | Human | 2012 |
| OP601603.1 | PB1 | segment_2 | Avian | 2019 |
| OP221316.1 | PB1 | segment_2 | Avian | 2022 |
| PV410879.1 | PB1 | segment_2 | Avian | 2021 |
| PV421709.1 | PB1 | segment_2 | Avian | 2022 |
| EU871834.2 | PB1 | segment_2 | Avian | 1987 |
| OL370085.1 | PB1 | segment_2 | Avian | 2015 |
| CY077691.1 | PB1 | segment_2 | Avian | 1981 |
| KJ729349.1 | PB1 | segment_2 | Avian | 2013 |
| AF389116.1 | PB1 | segment_2 | Human | 1934 |
| OP535289.1 | PB1 | segment_2 | Avian | 2022 |
| MH499234.1 | PB1 | segment_2 | Environment | 2016 |
| MH560232.1 | PB1 | segment_2 | Avian | 2014 |
| MK996234.1 | PB1 | segment_2 | Avian | 2019 |
| MN934004.1 | PB1 | segment_2 | Non-human mammal | 2019 |
| OQ780920.1 | PB1 | segment_2 | Avian | 2018 |
| EU798909.1 | PB1 | segment_2 | Non-human mammal | 2004 |
| PQ689842.1 | PB1 | segment_2 | Human | 2020 |
| CY028802.1 | PB1 | segment_2 | Non-human mammal | 1978 |
| OQ426554.1 | PB1 | segment_2 | Non-human mammal | 2021 |
| JN869515.1 | PB1 | segment_2 | Avian | 2009 |
| KF260579.1 | PB1 | segment_2 | Avian | 2007 |
| MW295731.1 | PB1 | segment_2 | Non-human mammal | 2014 |
| PP274818.1 | PB1 | segment_2 | Non-human mammal | 2023 |
| PP345552.1 | PB1 | segment_2 | Avian | 2019 |
| PQ191260.1 | PB1 | segment_2 | Avian | 2023 |
| PQ327687.1 | PB1 | segment_2 | Avian | 2024 |
| CY089801.1 | PB1 | segment_2 | Non-human mammal | 2010 |
| JQ924795.1 | PB1 | segment_2 | Avian | 2011 |
| CY021059.1 | PB1 | segment_2 | Human | 1954 |
| PQ116667.1 | PB1 | segment_2 | Non-human mammal | 2022 |
| KR700646.1 | PB1 | segment_2 | Non-human mammal | 2009 |
| MN882447.1 | PB1 | segment_2 | Avian | 2018 |
| KJ493634.1 | PB1 | segment_2 | Non-human mammal | 2013 |
| PX457968.1 | PB1 | segment_2 | Avian | 2022 |
| AB434414.1 | PB1 | segment_2 | Non-human mammal | 1996 |
| OQ360582.1 | PB1 | segment_2 | Non-human mammal | 2022 |
| PV388399.1 | PB1 | segment_2 | Avian | 2024 |

| | | | | |
|---|---|---|---|---|
| HQ877025.1 | PB1 | segment_2 | Non-human mammal | 2009 |
| JN043460.1 | PB1 | segment_2 | Non-human mammal | 2008 |
| PQ101255.1 | PB1 | segment_2 | Human | 2024 |
| CY039397.1 | PB1 | segment_2 | Non-human mammal | 1966 |
| PV739619.1 | PB1 | segment_2 | Avian | 2025 |
| MW362660.1 | PB1 | segment_2 | Non-human mammal | 2015 |
| MH376908.1 | PB1 | segment_2 | Bat | 2017 |
| KF881681.1 | PB1 | segment_2 | Avian | 2013 |
| CY103874.1 | PB1 | segment_2 | Bat | 2009 |
| CY125943.1 | PB1 | segment_2 | Bat | 2010 |
| MH682201.1 | PB1 | segment_2 | Bat | 2012 |
| PV233912.1 | PB1 | segment_2 | Human | 2023 |
| OL355042.1 | PA | segment_3_PA | Avian | 2019 |
| CY240824.1 | PA | segment_3_PA | Avian | 2016 |
| CY005769.1 | PA | segment_3_PA | Avian | 1983 |
| GU052008.1 | PA | segment_3_PA | Avian | 1975 |
| CY004004.1 | PA | segment_3_PA | Avian | 1978 |
| CY004111.1 | PA | segment_3_PA | Avian | 1979 |
| MH592235.1 | PA | segment_3_PA | Avian | 2012 |
| CY014654.1 | PA | segment_3_PA | Avian | 1965 |
| CY003884.1 | PA | segment_3_PA | Avian | 1985 |
| CY003973.1 | PA | segment_3_PA | Avian | 1998 |
| KF435048.1 | PA | segment_3_PA | Avian | 1953 |
| GU122111.1 | PA | segment_3_PA | Avian | 2008 |
| CY005817.1 | PA | segment_3_PA | Avian | 1963 |
| CY240904.1 | PA | segment_3_PA | Environment | 2016 |
| KJ729350.1 | PA | segment_3_PA | Avian | 2013 |
| CY005820.1 | PA | segment_3_PA | Avian | 1956 |
| CY014684.1 | PA | segment_3_PA | Avian | 1956 |
| KY463821.1 | PA | segment_3_PA | Avian | 2014 |
| CY014676.1 | PA | segment_3_PA | Avian | 1949 |
| PV946097.1 | PA | segment_3_PA | Avian | 2024 |
| KU042230.1 | PA | segment_3_PA | Avian | 2013 |
| EU871833.1 | PA | segment_3_PA | Avian | 1987 |
| PV123271.1 | PA | segment_3_PA | Avian | 2024 |
| CY014996.1 | PA | segment_3_PA | Avian | 1927 |
| KF619246.1 | PA | segment_3_PA | Avian | 1925 |
| MZ410775.1 | PA | segment_3_PA | Avian | 2020 |
| CY028841.1 | PA | segment_3_PA | Non-human mammal | 1963 |
| CY005863.1 | PA | segment_3_PA | Avian | 1980 |
| PV554096.1 | PA | segment_3_PA | MISSING | 2023 |

| | | | | |
|---|---|---|---|---|
| GU052318.1 | PA | segment_3_PA | Avian | 1992 |
| OQ861209.1 | PA | segment_3_PA | Avian | 2022 |
| CY015051.1 | PA | segment_3_PA | Avian | 1934 |
| HQ221596.1 | PA | segment_3_PA | Avian | 2009 |
| MW107383.1 | PA | segment_3_PA | Avian | 2017 |
| PV421859.1 | PA | segment_3_PA | Avian | 2023 |
| PP929865.1 | PA | segment_3_PA | Avian | 2024 |
| KY614998.1 | PA | segment_3_PA | Avian | 2016 |
| PV410190.1 | PA | segment_3_PA | Avian | 2021 |
| PV410198.1 | PA | segment_3_PA | Avian | 2022 |
| PQ665460.1 | PA | segment_3_PA | Human | 2024 |
| EU743056.1 | PA | segment_3_PA | Avian | 1987 |
| KF768218.1 | PA | segment_3_PA | Avian | 2006 |
| AF389117.1 | PA | segment_3_PA | Human | 1934 |
| KX523693.1 | PA | segment_3_PA | Avian | 2016 |
| KF768198.1 | PA | segment_3_PA | Avian | 2008 |
| MN882432.1 | PA | segment_3_PA | Avian | 2017 |
| JX273137.1 | PA | segment_3_PA | Avian | 2011 |
| KU042740.1 | PA | segment_3_PA | Avian | 2014 |
| MW356860.1 | PA | segment_3_PA | Avian | 2015 |
| OQ367231.1 | PA | segment_3_PA | Human | 2022 |
| HQ877026.1 | PA | segment_3_PA | Non-human mammal | 2009 |
| MZ945785.1 | PA | segment_3_PA | Non-human mammal | 2019 |
| KJ162619.1 | PA | segment_3_PA | Avian | 2013 |
| OP535170.1 | PA | segment_3_PA | Avian | 2022 |
| PQ356533.1 | PA | segment_3_PA | Human | 2024 |
| PV875053.1 | PA | segment_3_PA | Avian | 2024 |
| MW244134.1 | PA | segment_3_PA | Non-human mammal | 2020 |
| PV949366.1 | PA | segment_3_PA | Avian | 2024 |
| CY085837.2 | PA | segment_3_PA | Non-human mammal | 2007 |
| PX457969.1 | PA | segment_3_PA | Avian | 2022 |
| PP338283.1 | PA | segment_3_PA | Non-human mammal | 2021 |
| OQ836215.1 | PA | segment_3_PA | Avian | 2019 |
| AB434415.1 | PA | segment_3_PA | Non-human mammal | 1996 |
| CY039396.1 | PA | segment_3_PA | Non-human mammal | 1966 |
| PV875061.1 | PA | segment_3_PA | Avian | 2023 |
| MZ073289.1 | PA | segment_3_PA | Bat | 2018 |
| MH376907.1 | PA | segment_3_PA | Bat | 2017 |
| KJ493635.1 | PA | segment_3_PA | Non-human mammal | 2013 |
| OQ459925.1 | PA | segment_3_PA | Non-human mammal | 2021 |
| FJ844363.1 | PA | segment_3_PA | Non-human mammal | 2005 |

| | | | | |
|---|---|---|---|---|
| OR569040.1 | PA | segment_3_PA | Avian | 2020 |
| MH682202.1 | PA | segment_3_PA | Bat | 2012 |
| CY125944.1 | PA | segment_3_PA | Bat | 2010 |
| CY103875.1 | PA | segment_3_PA | Bat | 2009 |
| OM585572.1 | PA | segment_3_PA | Human | 2021 |
| OP292921.1 | HA | segment_4 | Non-human mammal | 2022 |
| PV233914.1 | HA | segment_4 | Human | 2023 |
| OQ216561.1 | HA | segment_4 | Bat | 2018 |
| MH764237.1 | HA | segment_4 | Avian | 2017 |
| CY103876.1 | HA | segment_4 | Bat | 2009 |
| LC728361.1 | HA | segment_4 | Non-human mammal | 2022 |
| PV133383.1 | HA | segment_4 | Avian | 2025 |
| OL371919.1 | HA | segment_4 | Environment | 2019 |
| MK928220.1 | HA | segment_4 | Avian | 2016 |
| CY116759.1 | HA | segment_4 | Avian | 1981 |
| CY014710.1 | HA | segment_4 | Avian | 1973 |
| CY014976.1 | HA | segment_4 | Human | 1957 |
| AF290439.1 | HA | segment_4 | Avian | 1976 |
| CY094365.1 | HA | segment_4 | Avian | 2007 |
| CY003847.1 | HA | segment_4 | Avian | 1977 |
| CY003871.1 | HA | segment_4 | Avian | 1984 |
| MT624466.1 | HA | segment_4 | Avian | 2019 |
| KY272859.1 | HA | segment_4 | Avian | 2016 |
| MZ575090.1 | HA | segment_4 | Avian | 2014 |
| CY005808.1 | HA | segment_4 | Avian | 1977 |
| CY042290.1 | HA | segment_4 | Avian | 1997 |
| CY125854.1 | HA | segment_4 | Human | 1961 |
| MH498654.1 | HA | segment_4 | Environment | 2016 |
| OR037484.1 | HA | segment_4 | Avian | 2021 |
| CY036871.1 | HA | segment_4 | Non-human mammal | 1975 |
| OP535291.1 | HA | segment_4 | Avian | 2022 |
| AB558258.1 | HA | segment_4 | Avian | 2002 |
| KP766054.1 | HA | segment_4 | Avian | 2014 |
| CY006009.1 | HA | segment_4 | Avian | 1983 |
| MW310457.1 | HA | segment_4 | Avian | 2008 |
| AF194990.1 | HA | segment_4 | Avian | 1997 |
| PP864033.1 | HA | segment_4 | Non-human mammal | 2023 |
| OP435644.1 | HA | segment_4 | Human | 2019 |
| PV738497.1 | HA | segment_4 | Non-human mammal | 2025 |
| DQ376621.1 | HA | segment_4 | Avian | 1997 |
| CY130030.1 | HA | segment_4 | Avian | 1965 |

| | | | | |
|---|---|---|---|---|
| CY125945.1 | HA | segment_4 | Bat | 2010 |
| LC497097.1 | HA | segment_4 | Avian | 2018 |
| MN530432.1 | HA | segment_4 | Environment | 2007 |
| AY968676.1 | HA | segment_4 | Avian | 1963 |
| CY004036.1 | HA | segment_4 | Avian | 1978 |
| CY004054.1 | HA | segment_4 | Avian | 1978 |
| CY110921.1 | HA | segment_4 | Human | 1934 |
| OQ737469.1 | HA | segment_4 | Non-human mammal | 2021 |
| PP691660.1 | HA | segment_4 | Avian | 2023 |
| MW345940.1 | HA | segment_4 | Avian | 2019 |
| MN932186.1 | HA | segment_4 | Non-human mammal | 2018 |
| FJ750571.1 | HA | segment_4 | Avian | 1999 |
| AB542809.1 | HA | segment_4 | Avian | 1978 |
| AB558457.1 | HA | segment_4 | Avian | 1981 |
| CY034681.1 | HA | segment_4 | Avian | 1993 |
| CY110924.2 | HA | segment_4 | Human | 2009 |
| PV337324.1 | HA | segment_4 | Avian | 2025 |
| KJ508885.1 | HA | segment_4 | Avian | 2010 |
| HM059998.1 | HA | segment_4 | Avian | 2006 |
| KX977730.1 | HA | segment_4 | Avian | 2008 |
| CY014636.1 | HA | segment_4 | Avian | 1981 |
| PP829826.1 | HA | segment_4 | Environment | 2023 |
| MN911205.1 | HA | segment_4 | Avian | 2019 |
| MG214160.1 | HA | segment_4 | Avian | 2017 |
| MG891872.2 | HA | segment_4 | Avian | 2017 |
| MH453821.1 | HA | segment_4 | Avian | 2014 |
| PP555669.1 | HA | segment_4 | Human | 2024 |
| PX129094.1 | HA | segment_4 | Avian | 2024 |
| FJ190111.1 | HA | segment_4 | Avian | 2001 |
| PP502447.1 | HA | segment_4 | Avian | 2019 |
| MN530511.1 | HA | segment_4 | Avian | 2016 |
| PV875062.1 | HA | segment_4 | Avian | 2023 |
| OL371391.1 | HA | segment_4 | Environment | 2019 |
| PP829645.1 | HA | segment_4 | Environment | 2023 |
| ON637239.1 | HA | segment_4 | Avian | 2008 |
| PV410207.1 | HA | segment_4 | Avian | 2022 |
| MW170290.1 | HA | segment_4 | Non-human mammal | 2019 |
| KP186856.1 | HA | segment_4 | Avian | 2013 |
| KP186859.1 | HA | segment_4 | Avian | 2013 |
| PV233909.1 | NP | segment_5 | Human | 2023 |
| PV449020.1 | NP | segment_5 | Non-human mammal | 2024 |

| | | | | |
|---|---|---|---|---|
| AB914485.1 | NP | segment_5 | Non-human mammal | 2012 |
| PX271017.1 | NP | segment_5 | Avian | 2024 |
| HM193673.1 | NP | segment_5 | Avian | 2006 |
| GU053298.1 | NP | segment_5 | Avian | 2004 |
| AY611527.1 | NP | segment_5 | Avian | 2004 |
| EU735837.2 | NP | segment_5 | Avian | 1995 |
| AY633215.1 | NP | segment_5 | Avian | 1998 |
| CY004292.1 | NP | segment_5 | Avian | 1976 |
| CY014824.1 | NP | segment_5 | Avian | 1995 |
| CY003955.1 | NP | segment_5 | Avian | 1996 |
| EU557532.1 | NP | segment_5 | Avian | 2006 |
| MW099136.1 | NP | segment_5 | Avian | 2017 |
| CY177521.1 | NP | segment_5 | Avian | 1976 |
| MT261595.1 | NP | segment_5 | Avian | 2019 |
| CY004793.1 | NP | segment_5 | Avian | 1979 |
| CY004229.1 | NP | segment_5 | Avian | 1990 |
| AY633119.1 | NP | segment_5 | Avian | 1983 |
| GU052002.1 | NP | segment_5 | Avian | 1975 |
| CY093761.1 | NP | segment_5 | Avian | 2008 |
| CY094376.1 | NP | segment_5 | Avian | 2007 |
| CY181340.1 | NP | segment_5 | Avian | 1974 |
| GU052316.2 | NP | segment_5 | Avian | 1992 |
| AB932557.1 | NP | segment_5 | Avian | 2014 |
| CY015076.1 | NP | segment_5 | Avian | 1983 |
| EU182261.2 | NP | segment_5 | Avian | 2001 |
| KF042075.1 | NP | segment_5 | Avian | 2013 |
| CY005259.1 | NP | segment_5 | Avian | 2004 |
| PP474947.1 | NP | segment_5 | Avian | 2023 |
| MW727369.1 | NP | segment_5 | Avian | 2018 |
| MH170290.1 | NP | segment_5 | Avian | 2016 |
| MG366513.1 | NP | segment_5 | Avian | 2016 |
| MK339598.1 | NP | segment_5 | Non-human mammal | 2012 |
| KU860474.1 | NP | segment_5 | Avian | 1980 |
| CY014994.1 | NP | segment_5 | Avian | 1927 |
| CY004446.1 | NP | segment_5 | Avian | 2004 |
| LC486290.1 | NP | segment_5 | Avian | 2017 |
| MF510853.1 | NP | segment_5 | Avian | 2017 |
| PP829811.1 | NP | segment_5 | Environment | 2023 |
| KR824583.1 | NP | segment_5 | Avian | 2010 |
| LC497130.1 | NP | segment_5 | Environment | 2018 |
| KX979010.1 | NP | segment_5 | Avian | 2009 |

| | | | | |
|---|---|---|---|---|
| PQ115509.1 | NP | segment_5 | Non-human mammal | 2020 |
| OQ168180.1 | NP | segment_5 | Non-human mammal | 2019 |
| OL371552.1 | NP | segment_5 | Environment | 2019 |
| MN530615.1 | NP | segment_5 | Avian | 2016 |
| OL370293.1 | NP | segment_5 | Environment | 2018 |
| CY029892.1 | NP | segment_5 | Avian | 2004 |
| PV387981.1 | NP | segment_5 | Avian | 2024 |
| PV123273.1 | NP | segment_5 | Avian | 2024 |
| ON911304.1 | NP | segment_5 | Avian | 2021 |
| HM145068.1 | NP | segment_5 | Avian | 2004 |
| EU871831.1 | NP | segment_5 | Avian | 1987 |
| OP458635.1 | NP | segment_5 | Non-human mammal | 2020 |
| MG280144.1 | NP | segment_5 | Avian | 2016 |
| CY005803.1 | NP | segment_5 | Non-human mammal | 1956 |
| HQ877028.1 | NP | segment_5 | Non-human mammal | 2009 |
| CY005416.1 | NP | segment_5 | Avian | 1990 |
| CY246254.1 | NP | segment_5 | Non-human mammal | 2016 |
| OQ367072.1 | NP | segment_5 | Human | 2022 |
| PX324940.1 | NP | segment_5 | Human | 2019 |
| KP657910.1 | NP | segment_5 | Avian | 2014 |
| HM142749.1 | NP | segment_5 | Non-human mammal | 2005 |
| KU860468.1 | NP | segment_5 | Avian | 1980 |
| MN418734.1 | NP | segment_5 | Non-human mammal | 2011 |
| MH991780.1 | NP | segment_5 | Avian | 2014 |
| MG962433.1 | NP | segment_5 | Non-human mammal | 2012 |
| AF258516.1 | NP | segment_5 | Environment | 1998 |
| PX460739.1 | NP | segment_5 | Avian | 2023 |
| MZ073287.1 | NP | segment_5 | Bat | 2018 |
| KX344897.1 | NP | segment_5 | Avian | 1980 |
| MN418736.1 | NP | segment_5 | Non-human mammal | 2012 |
| CY125946.1 | NP | segment_5 | Bat | 2010 |
| CY103877.1 | NP | segment_5 | Bat | 2009 |
| MH453833.1 | NA | segment_6 | Avian | 2013 |
| KY644191.1 | NA | segment_6 | Avian | 2016 |
| PV770036.1 | NA | segment_6 | Human | 2024 |
| PX457169.1 | NA | segment_6 | Avian | 2016 |
| OL370288.1 | NA | segment_6 | Environment | 2018 |
| CY213697.1 | NA | segment_6 | Avian | 2015 |
| PX063872.1 | NA | segment_6 | Avian | 2024 |
| EU871830.1 | NA | segment_6 | Avian | 1987 |
| KY013750.1 | NA | segment_6 | Avian | 2015 |

| | | | | |
|---|---|---|---|---|
| MZ364006.1 | NA | segment_6 | Non-human mammal | 2018 |
| MW874986.1 | NA | segment_6 | Avian | 2020 |
| MH347051.1 | NA | segment_6 | Environment | 2017 |
| MH134851.1 | NA | segment_6 | Avian | 2016 |
| CY005853.1 | NA | segment_6 | Avian | 1999 |
| PP502591.1 | NA | segment_6 | Avian | 2017 |
| PV412289.1 | NA | segment_6 | Human | 2023 |
| PQ675398.1 | NA | segment_6 | Avian | 2024 |
| CY036873.1 | NA | segment_6 | Non-human mammal | 1975 |
| MH498747.1 | NA | segment_6 | Avian | 2016 |
| KF435051.1 | NA | segment_6 | Avian | 1953 |
| MK629014.1 | NA | segment_6 | Avian | 2018 |
| PP549229.1 | NA | segment_6 | Avian | 2022 |
| PX312471.1 | NA | segment_6 | Avian | 2024 |
| CY014993.1 | NA | segment_6 | Avian | 1927 |
| PP664081.1 | NA | segment_6 | Environment | 2022 |
| KP766161.1 | NA | segment_6 | Avian | 2014 |
| GU122057.1 | NA | segment_6 | Avian | 2007 |
| MZ073285.1 | NA | segment_6 | Bat | 2018 |
| MN911195.1 | NA | segment_6 | Avian | 2019 |
| OP535173.1 | NA | segment_6 | Avian | 2022 |
| PV233908.1 | NA | segment_6 | Human | 2023 |
| MH682205.1 | NA | segment_6 | Bat | 2012 |
| KR077934.1 | NA | segment_6 | Bat | 2011 |
| KX185911.1 | NA | segment_6 | Avian | 2015 |
| KX571080.1 | NA | segment_6 | Non-human mammal | 2014 |
| CY103878.1 | NA | segment_6 | Bat | 2009 |
| CY103894.1 | NA | segment_6 | Bat | 2010 |
| OL371105.1 | NA | segment_6 | Avian | 2020 |
| OP430901.1 | NA | segment_6 | Human | 2019 |
| CY003986.1 | NA | segment_6 | Avian | 2002 |
| CY004717.1 | NA | segment_6 | Avian | 1977 |
| EU429793.1 | NA | segment_6 | Avian | 1968 |
| GQ176110.1 | NA | segment_6 | Avian | 1988 |
| CY014625.1 | NA | segment_6 | Avian | 1979 |
| GQ176118.1 | NA | segment_6 | Avian | 1985 |
| KY644153.1 | NA | segment_6 | Avian | 2015 |
| CY025255.1 | NA | segment_6 | Non-human mammal | 1987 |
| GQ161114.1 | NA | segment_6 | Non-human mammal | 2003 |
| CY115837.1 | NA | segment_6 | Non-human mammal | 2005 |
| CY115953.1 | NA | segment_6 | Non-human mammal | 2005 |

| | | | | |
|---|---|---|---|---|
| AB434306.1 | NA | segment_6 | Non-human mammal | 2004 |
| AB434298.1 | NA | segment_6 | Non-human mammal | 2003 |
| CY098502.1 | NA | segment_6 | Non-human mammal | 1994 |
| AB434290.1 | NA | segment_6 | Non-human mammal | 2000 |
| CY116398.1 | NA | segment_6 | Non-human mammal | 1995 |
| CY037962.1 | NA | segment_6 | Non-human mammal | 1999 |
| CY115875.1 | NA | segment_6 | Non-human mammal | 2001 |
| CY061639.1 | NA | segment_6 | Non-human mammal | 2009 |
| MK364857.1 | NA | segment_6 | Non-human mammal | 2004 |
| CY115906.1 | NA | segment_6 | Non-human mammal | 2007 |
| CY115929.1 | NA | segment_6 | Non-human mammal | 2003 |
| KR700681.1 | NA | segment_6 | Non-human mammal | 2011 |
| KC209522.1 | NA | segment_6 | Non-human mammal | 2012 |
| MZ410778.1 | NA | segment_6 | Avian | 2020 |
| PP333643.1 | NA | segment_6 | Avian | 2023 |
| AB472016.1 | NA | segment_6 | Avian | 2005 |
| PQ318337.1 | NA | segment_6 | Avian | 2023 |
| PX129124.1 | NA | segment_6 | Avian | 2025 |
| PV796172.1 | NA | segment_6 | Avian | 2022 |
| MN933417.1 | NA | segment_6 | Non-human mammal | 2018 |
| MN933410.1 | NA | segment_6 | Non-human mammal | 2019 |
| MW269564.1 | NA | segment_6 | Non-human mammal | 2017 |
| MT887311.1 | NA | segment_6 | Non-human mammal | 2020 |
| PV875064.1 | NA | segment_6 | Avian | 2023 |
| MH271630.1 | NA | segment_6 | Avian | 2017 |
| MH376903.1 | M1 | segment_7_M1 | Bat | 2017 |
| CY015109.1 | M1 | segment_7_M1 | Avian | 1983 |
| CY116399.1 | M1 | segment_7_M1 | Non-human mammal | 1995 |
| PQ358082.1 | M1 | segment_7_M1 | Avian | 2024 |
| AF231360.1 | M1 | segment_7_M1 | Avian | 1978 |
| AY300957.2 | M1 | segment_7_M1 | Avian | 2000 |
| CY005839.1 | M1 | segment_7_M1 | Avian | 1997 |
| CY003895.1 | M1 | segment_7_M1 | Avian | 1986 |
| CY004290.1 | M1 | segment_7_M1 | Avian | 1976 |
| CY003985.1 | M1 | segment_7_M1 | Avian | 2002 |
| CY015028.1 | M1 | segment_7_M1 | Avian | 2002 |
| DQ299830.1 | M1 | segment_7_M1 | Avian | 2004 |
| EU871829.2 | M1 | segment_7_M1 | Avian | 1987 |
| FJ752505.1 | M1 | segment_7_M1 | Non-human mammal | 2000 |
| CY005414.1 | M1 | segment_7_M1 | Avian | 1990 |
| AF231358.1 | M1 | segment_7_M1 | Environment | 1957 |

| | | | | |
|---|---|---|---|---|
| GU396852.1 | M1 | segment_7_M1 | Avian | 2006 |
| CY094254.1 | M1 | segment_7_M1 | Avian | 2008 |
| CY213552.1 | M1 | segment_7_M1 | Avian | 2015 |
| AB753188.1 | M1 | segment_7_M1 | Avian | 2007 |
| HE802742.1 | M1 | segment_7_M1 | Avian | 2006 |
| AB591848.2 | M1 | segment_7_M1 | Non-human mammal | 2007 |
| CY085425.1 | M1 | segment_7_M1 | Non-human mammal | 2001 |
| OQ861124.1 | M1 | segment_7_M1 | Avian | 2022 |
| AB285935.1 | M1 | segment_7_M1 | Human | 2001 |
| KP657912.1 | M1 | segment_7_M1 | Avian | 2014 |
| MW107875.1 | M1 | segment_7_M1 | Avian | 2017 |
| KJ162719.1 | M1 | segment_7_M1 | Avian | 2013 |
| CY004073.1 | M1 | segment_7_M1 | Avian | 1979 |
| MZ073286.1 | M1 | segment_7_M1 | Bat | 2018 |
| OL370690.1 | M1 | segment_7_M1 | Avian | 2017 |
| PP276148.1 | M1 | segment_7_M1 | Avian | 2022 |
| MG962435.1 | M1 | segment_7_M1 | Non-human mammal | 2012 |
| CY015047.1 | M1 | segment_7_M1 | Avian | 1934 |
| KX977916.1 | M1 | segment_7_M1 | Avian | 2007 |
| KM032356.1 | M1 | segment_7_M1 | Avian | 2009 |
| KT936660.1 | M1 | segment_7_M1 | Avian | 2013 |
| PV647681.1 | M1 | segment_7_M1 | Non-human mammal | 2024 |
| MW100599.1 | M1 | segment_7_M1 | Avian | 2017 |
| GU049952.1 | M1 | segment_7_M1 | Environment | 2007 |
| GU903111.1 | M1 | segment_7_M1 | Avian | 2008 |
| KY888127.1 | M1 | segment_7_M1 | Non-human mammal | 2016 |
| MZ410779.1 | M1 | segment_7_M1 | Avian | 2020 |
| CY029890.1 | M1 | segment_7_M1 | Avian | 2004 |
| ON366872.1 | M1 | segment_7_M1 | Avian | 2021 |
| JX096507.1 | M1 | segment_7_M1 | Non-human mammal | 2011 |
| KJ162591.1 | M1 | segment_7_M1 | Avian | 2012 |
| MT261389.1 | M1 | segment_7_M1 | Avian | 2019 |
| MT261401.1 | M1 | segment_7_M1 | Avian | 2019 |
| PV391116.1 | M1 | segment_7_M1 | Avian | 2024 |
| MW103579.1 | M1 | segment_7_M1 | Avian | 2016 |
| MN988776.1 | M1 | segment_7_M1 | Avian | 2019 |
| MW101290.1 | M1 | segment_7_M1 | Avian | 2017 |
| MW107355.1 | M1 | segment_7_M1 | Avian | 2016 |
| MW102899.1 | M1 | segment_7_M1 | Avian | 2017 |
| PV943724.1 | M1 | segment_7_M1 | Avian | 2025 |
| PP896388.1 | M1 | segment_7_M1 | Non-human mammal | 2022 |

| | | | | |
|---|---|---|---|---|
| MK117073.1 | M1 | segment_7_M1 | Human | 2017 |
| PQ115391.1 | M1 | segment_7_M1 | Non-human mammal | 2019 |
| KC167169.1 | M1 | segment_7_M1 | Non-human mammal | 2012 |
| MN933443.1 | M1 | segment_7_M1 | Non-human mammal | 2019 |
| CY098249.1 | M1 | segment_7_M1 | Avian | 2010 |
| OL354877.1 | M1 | segment_7_M1 | Avian | 2019 |
| GQ335501.1 | M1 | segment_7_M1 | Avian | 2006 |
| CY005801.1 | M1 | segment_7_M1 | Non-human mammal | 1956 |
| PQ070773.1 | M1 | segment_7_M1 | Human | 2024 |
| AF250491.1 | M1 | segment_7_M1 | Avian | 1997 |
| MW103115.1 | M1 | segment_7_M1 | Avian | 2017 |
| KC610052.1 | M1 | segment_7_M1 | Non-human mammal | 2010 |
| CY103879.1 | M1 | segment_7_M1 | Bat | 2009 |
| CY125948.1 | M1 | segment_7_M1 | Bat | 2010 |
| KR077935.1 | M1 | segment_7_M1 | Bat | 2011 |
| KU143704.1 | M1 | segment_7_M1 | Human | 2009 |
| KU143693.1 | M1 | segment_7_M1 | Human | 2009 |
| PV233915.1 | M1 | segment_7_M1 | Human | 2023 |
| KR233677.1 | M2 | segment_7_M2 | Avian | 2014 |
| CY048679.1 | M2 | segment_7_M2 | Avian | 2007 |
| AB304150.1 | M2 | segment_7_M2 | Avian | 2004 |
| MK886524.1 | M2 | segment_7_M2 | Non-human mammal | 2018 |
| CY103879.1 | M2 | segment_7_M2 | Bat | 2009 |
| CY125948.1 | M2 | segment_7_M2 | Bat | 2010 |
| KR077935.1 | M2 | segment_7_M2 | Bat | 2011 |
| OL355038.1 | M2 | segment_7_M2 | Environment | 2019 |
| GU050160.1 | M2 | segment_7_M2 | Avian | 2007 |
| AJ298948.1 | M2 | segment_7_M2 | Environment | 1969 |
| MK478680.1 | M2 | segment_7_M2 | Non-human mammal | 2019 |
| AB600868.1 | M2 | segment_7_M2 | Non-human mammal | 2005 |
| JN391251.1 | M2 | segment_7_M2 | Non-human mammal | 2004 |
| KM028938.1 | M2 | segment_7_M2 | Non-human mammal | 2011 |
| CY246653.1 | M2 | segment_7_M2 | Non-human mammal | 2016 |
| KY971199.1 | M2 | segment_7_M2 | Avian | 2013 |
| KM892801.1 | M2 | segment_7_M2 | Avian | 2009 |
| PV653848.1 | M2 | segment_7_M2 | Avian | 2024 |
| MW621567.1 | M2 | segment_7_M2 | Non-human mammal | 2020 |
| KM244111.1 | M2 | segment_7_M2 | Avian | 2014 |
| KJ729346.1 | M2 | segment_7_M2 | Avian | 2007 |
| AB847432.1 | M2 | segment_7_M2 | Avian | 2012 |
| CY005839.1 | M2 | segment_7_M2 | Avian | 1997 |

| | | | | |
|---|---|---|---|---|
| MW110059.1 | M2 | segment_7_M2 | Avian | 2016 |
| AB753188.1 | M2 | segment_7_M2 | Avian | 2007 |
| CY005445.1 | M2 | segment_7_M2 | Avian | 2001 |
| CY003895.1 | M2 | segment_7_M2 | Avian | 1986 |
| CY004073.1 | M2 | segment_7_M2 | Avian | 1979 |
| MN049529.1 | M2 | segment_7_M2 | Avian | 2018 |
| OM527224.1 | M2 | segment_7_M2 | Avian | 2021 |
| MN049536.1 | M2 | segment_7_M2 | Avian | 2018 |
| CY032318.1 | M2 | segment_7_M2 | Non-human mammal | 1989 |
| JN391235.1 | M2 | segment_7_M2 | Non-human mammal | 2003 |
| CY085497.2 | M2 | segment_7_M2 | Non-human mammal | 2002 |
| KM366696.1 | M2 | segment_7_M2 | Human | 2014 |
| JN043435.1 | M2 | segment_7_M2 | Non-human mammal | 2009 |
| KP657912.1 | M2 | segment_7_M2 | Avian | 2014 |
| MW102835.1 | M2 | segment_7_M2 | Avian | 2017 |
| PP274916.1 | M2 | segment_7_M2 | Non-human mammal | 2023 |
| AB775609.1 | M2 | segment_7_M2 | Avian | 2012 |
| OP535174.1 | M2 | segment_7_M2 | Avian | 2022 |
| EU662948.1 | M2 | segment_7_M2 | Avian | 2004 |
| MN882404.1 | M2 | segment_7_M2 | Avian | 2017 |
| MN882508.1 | M2 | segment_7_M2 | Avian | 2019 |
| KC859057.1 | M2 | segment_7_M2 | Non-human mammal | 2010 |
| MN267533.1 | M2 | segment_7_M2 | Avian | 2012 |
| OR664498.1 | M2 | segment_7_M2 | Avian | 2023 |
| GQ373129.1 | M2 | segment_7_M2 | Avian | 2007 |
| JN200413.1 | M2 | segment_7_M2 | Avian | 2010 |
| GU122045.1 | M2 | segment_7_M2 | Non-human mammal | 2008 |
| OR473200.1 | M2 | segment_7_M2 | Non-human mammal | 2018 |
| KF638582.1 | M2 | segment_7_M2 | Avian | 2012 |
| MN933445.1 | M2 | segment_7_M2 | Non-human mammal | 2019 |
| ON368086.1 | M2 | segment_7_M2 | Avian | 2020 |
| MG280628.1 | M2 | segment_7_M2 | Avian | 2011 |
| MN933455.1 | M2 | segment_7_M2 | Non-human mammal | 2018 |
| KC610021.1 | M2 | segment_7_M2 | Non-human mammal | 2011 |
| AB600865.1 | M2 | segment_7_M2 | Non-human mammal | 2004 |
| CY089799.1 | M2 | segment_7_M2 | Non-human mammal | 2010 |
| KU143688.1 | M2 | segment_7_M2 | Human | 2009 |
| MH168318.1 | M2 | segment_7_M2 | Human | 2015 |
| CY245658.1 | M2 | segment_7_M2 | Non-human mammal | 2015 |
| JX499146.1 | M2 | segment_7_M2 | Avian | 2009 |
| OQ835506.1 | M2 | segment_7_M2 | Avian | 2022 |

| | | | | |
|---|---|---|---|---|
| MN169178.1 | M2 | segment_7_M2 | Human | 2019 |
| MG962475.1 | M2 | segment_7_M2 | Non-human mammal | 2012 |
| AB285938.1 | M2 | segment_7_M2 | Human | 2001 |
| AB795909.1 | M2 | segment_7_M2 | Environment | 2008 |
| CY115999.1 | M2 | segment_7_M2 | Non-human mammal | 1987 |
| JQ396191.2 | M2 | segment_7_M2 | Human | 2010 |
| PV422111.1 | M2 | segment_7_M2 | Avian | 2022 |
| CY005801.1 | M2 | segment_7_M2 | Non-human mammal | 1956 |
| MZ073286.1 | M2 | segment_7_M2 | Bat | 2018 |
| MH376903.1 | M2 | segment_7_M2 | Bat | 2017 |
| DQ525415.1 | M2 | segment_7_M2 | Avian | 2001 |
| MH682207.1 | NS1 | segment_8_NS1 | Bat | 2012 |
| CY125949.1 | NS1 | segment_8_NS1 | Bat | 2010 |
| CY103880.1 | NS1 | segment_8_NS1 | Bat | 2009 |
| MZ564696.1 | NS1 | segment_8_NS1 | Avian | 2019 |
| OR044200.1 | NS1 | segment_8_NS1 | Environment | 2020 |
| OQ120654.1 | NS1 | segment_8_NS1 | Avian | 2015 |
| PP829831.1 | NS1 | segment_8_NS1 | Environment | 2023 |
| AB781665.1 | NS1 | segment_8_NS1 | Avian | 2011 |
| MH578809.1 | NS1 | segment_8_NS1 | Avian | 2005 |
| PQ031950.1 | NS1 | segment_8_NS1 | Avian | 2023 |
| PQ191266.1 | NS1 | segment_8_NS1 | Avian | 2023 |
| OQ413791.1 | NS1 | segment_8_NS1 | Avian | 2022 |
| MF046319.2 | NS1 | segment_8_NS1 | Avian | 2016 |
| OQ793889.1 | NS1 | segment_8_NS1 | Avian | 2017 |
| OP806456.1 | NS1 | segment_8_NS1 | Avian | 2022 |
| CY093092.1 | NS1 | segment_8_NS1 | Avian | 2010 |
| MN882453.1 | NS1 | segment_8_NS1 | Avian | 2018 |
| PQ032005.1 | NS1 | segment_8_NS1 | Avian | 2023 |
| CY109814.1 | NS1 | segment_8_NS1 | Avian | 2007 |
| GU051990.1 | NS1 | segment_8_NS1 | Avian | 2007 |
| FJ205646.1 | NS1 | segment_8_NS1 | Avian | 1998 |
| KY561224.1 | NS1 | segment_8_NS1 | Avian | 2015 |
| DQ997550.1 | NS1 | segment_8_NS1 | Avian | 2003 |
| JN244253.1 | NS1 | segment_8_NS1 | Avian | 2010 |
| OL370334.1 | NS1 | segment_8_NS1 | Environment | 2019 |
| KU320891.1 | NS1 | segment_8_NS1 | Avian | 2015 |
| EU249549.1 | NS1 | segment_8_NS1 | Avian | 2004 |
| PV315204.1 | NS1 | segment_8_NS1 | Avian | 2022 |
| ON716358.1 | NS1 | segment_8_NS1 | Avian | 2021 |
| MK213335.1 | NS1 | segment_8_NS1 | Avian | 2017 |

| | | | | |
|---|---|---|---|---|
| OL370260.1 | NS1 | segment_8_NS1 | Avian | 2017 |
| AB775626.1 | NS1 | segment_8_NS1 | Avian | 2012 |
| MW333194.1 | NS1 | segment_8_NS1 | Avian | 2017 |
| OR267222.1 | NS1 | segment_8_NS1 | Avian | 2022 |
| MG241347.1 | NS1 | segment_8_NS1 | Avian | 2017 |
| LC486293.1 | NS1 | segment_8_NS1 | Avian | 2017 |
| AB275410.1 | NS1 | segment_8_NS1 | Avian | 2004 |
| AF250495.1 | NS1 | segment_8_NS1 | Avian | 1997 |
| KX518665.1 | NS1 | segment_8_NS1 | Avian | 2014 |
| GU050283.1 | NS1 | segment_8_NS1 | Avian | 2006 |
| AB932560.1 | NS1 | segment_8_NS1 | Avian | 2014 |
| KX228214.1 | NS1 | segment_8_NS1 | Avian | 2014 |
| CY029255.1 | NS1 | segment_8_NS1 | Avian | 2003 |
| KJ162720.1 | NS1 | segment_8_NS1 | Avian | 2013 |
| OP535295.1 | NS1 | segment_8_NS1 | Avian | 2022 |
| OL450392.1 | NS1 | segment_8_NS1 | Environment | 2015 |
| MW101805.1 | NS1 | segment_8_NS1 | Avian | 2017 |
| LC364325.1 | NS1 | segment_8_NS1 | Avian | 2017 |
| PQ107775.1 | NS1 | segment_8_NS1 | Non-human mammal | 2022 |
| MZ477716.1 | NS1 | segment_8_NS1 | Non-human mammal | 2020 |
| MT666836.1 | NS1 | segment_8_NS1 | Non-human mammal | 2015 |
| MN939499.1 | NS1 | segment_8_NS1 | Non-human mammal | 2017 |
| AB795922.1 | NS1 | segment_8_NS1 | Environment | 2008 |
| CY116000.1 | NS1 | segment_8_NS1 | Non-human mammal | 1987 |
| AB434340.1 | NS1 | segment_8_NS1 | Non-human mammal | 2005 |
| PX101778.1 | NS1 | segment_8_NS1 | Human | 2021 |
| PP274917.1 | NS1 | segment_8_NS1 | Non-human mammal | 2023 |
| PX328778.1 | NS1 | segment_8_NS1 | Human | 2022 |
| PX335963.1 | NS1 | segment_8_NS1 | Human | 2019 |
| MN074254.1 | NS1 | segment_8_NS1 | Human | 2019 |
| MK300672.1 | NS1 | segment_8_NS1 | Non-human mammal | 2017 |
| LC430964.1 | NS1 | segment_8_NS1 | Non-human mammal | 2003 |
| JN043450.1 | NS1 | segment_8_NS1 | Non-human mammal | 2008 |
| PX444236.1 | NS1 | segment_8_NS1 | Non-human mammal | 2019 |
| MZ073288.1 | NS1 | segment_8_NS1 | Bat | 2018 |
| PQ115423.1 | NS1 | segment_8_NS1 | Non-human mammal | 2019 |
| MN939508.1 | NS1 | segment_8_NS1 | Non-human mammal | 2016 |
| OQ356153.1 | NS1 | segment_8_NS1 | Non-human mammal | 2022 |
| PQ115512.1 | NS1 | segment_8_NS1 | Non-human mammal | 2020 |
| MG962436.1 | NS1 | segment_8_NS1 | Non-human mammal | 2012 |
| PX444276.1 | NS1 | segment_8_NS1 | Non-human mammal | 2021 |

| | | | | |
|---|---|---|---|---|
| PQ894542.1 | NS1 | segment_8_NS1 | Avian | 2024 |
| CY005804.1 | NS1 | segment_8_NS1 | Non-human mammal | 1956 |
| LC415038.1 | NS1 | segment_8_NS1 | Avian | 2016 |
| PV233910.1 | NS1 | segment_8_NS1 | Human | 2023 |
| LC335987.1 | NEP | segment_8_NS2 | Avian | 2017 |
| DQ525413.1 | NEP | segment_8_NS2 | Avian | 2001 |
| EU599307.1 | NEP | segment_8_NS2 | Avian | 2005 |
| AB937725.1 | NEP | segment_8_NS2 | Avian | 2013 |
| LC106108.1 | NEP | segment_8_NS2 | Avian | 2015 |
| AB166866.1 | NEP | segment_8_NS2 | Avian | 2004 |
| AB786680.1 | NEP | segment_8_NS2 | Avian | 2012 |
| LC000600.1 | NEP | segment_8_NS2 | Avian | 2014 |
| AB735627.1 | NEP | segment_8_NS2 | Avian | 2012 |
| AB981454.1 | NEP | segment_8_NS2 | Avian | 2013 |
| EU743310.1 | NEP | segment_8_NS2 | Avian | 1986 |
| AB753181.1 | NEP | segment_8_NS2 | Avian | 2006 |
| AB753189.1 | NEP | segment_8_NS2 | Avian | 2007 |
| AB775803.1 | NEP | segment_8_NS2 | Avian | 2012 |
| CY003918.1 | NEP | segment_8_NS2 | Avian | 1988 |
| CY015077.1 | NEP | segment_8_NS2 | Avian | 1983 |
| CY005701.1 | NEP | segment_8_NS2 | Avian | 1980 |
| CY005417.1 | NEP | segment_8_NS2 | Avian | 1990 |
| AB735611.1 | NEP | segment_8_NS2 | Avian | 2012 |
| CY005404.1 | NEP | segment_8_NS2 | Avian | 1983 |
| AB593435.1 | NEP | segment_8_NS2 | Avian | 2006 |
| CY029539.1 | NEP | segment_8_NS2 | Avian | 2006 |
| AB727570.1 | NEP | segment_8_NS2 | Avian | 2012 |
| AB818501.1 | NEP | segment_8_NS2 | Avian | 2012 |
| EU301358.1 | NEP | segment_8_NS2 | Avian | 2005 |
| DQ299857.1 | NEP | segment_8_NS2 | Avian | 2004 |
| AB591849.2 | NEP | segment_8_NS2 | Non-human mammal | 2007 |
| CY032297.1 | NEP | segment_8_NS2 | Non-human mammal | 1963 |
| EU871832.1 | NEP | segment_8_NS2 | Avian | 1987 |
| CY003898.1 | NEP | segment_8_NS2 | Avian | 1986 |
| AB753212.1 | NEP | segment_8_NS2 | Avian | 2009 |
| AJ298949.1 | NEP | segment_8_NS2 | Environment | 1934 |
| CY011000.2 | NEP | segment_8_NS2 | Human | 2000 |
| AB442000.1 | NEP | segment_8_NS2 | Human | 2006 |
| CY020449.1 | NEP | segment_8_NS2 | Human | 1936 |
| AB514943.1 | NEP | segment_8_NS2 | Non-human mammal | 2008 |
| AB434292.1 | NEP | segment_8_NS2 | Non-human mammal | 2000 |

| | | | | |
|---|---|---|---|---|
| AB909402.1 | NEP | segment_8_NS2 | Non-human mammal | 2011 |
| AB434348.1 | NEP | segment_8_NS2 | Non-human mammal | 2003 |
| EU301336.1 | NEP | segment_8_NS2 | Non-human mammal | 2004 |
| EU409952.1 | NEP | segment_8_NS2 | Non-human mammal | 2007 |
| EU258935.1 | NEP | segment_8_NS2 | Non-human mammal | 2006 |
| AB434308.1 | NEP | segment_8_NS2 | Non-human mammal | 2004 |
| AB434340.1 | NEP | segment_8_NS2 | Non-human mammal | 2005 |
| AB434412.1 | NEP | segment_8_NS2 | Non-human mammal | 1980 |
| EU518716.1 | NEP | segment_8_NS2 | Avian | 2005 |
| CY005804.1 | NEP | segment_8_NS2 | Non-human mammal | 1956 |
| AJ298950.1 | NEP | segment_8_NS2 | Environment | 1983 |
| AB261857.1 | NEP | segment_8_NS2 | Avian | 2005 |
| CY005738.1 | NEP | segment_8_NS2 | Avian | 1976 |
| AY241629.1 | NEP | segment_8_NS2 | Avian | 2001 |
| AB916655.1 | NEP | segment_8_NS2 | Avian | 2013 |
| CY015030.1 | NEP | segment_8_NS2 | Avian | 2002 |
| CY005240.1 | NEP | segment_8_NS2 | Avian | 2000 |
| AB781665.1 | NEP | segment_8_NS2 | Avian | 2011 |
| AB275287.1 | NEP | segment_8_NS2 | Avian | 1980 |
| EU182269.3 | NEP | segment_8_NS2 | Avian | 2004 |
| LC000616.1 | NEP | segment_8_NS2 | Avian | 2013 |
| EU430506.1 | NEP | segment_8_NS2 | Avian | 2005 |
| CY103880.1 | NEP | segment_8_NS2 | Bat | 2009 |
| CY125949.1 | NEP | segment_8_NS2 | Bat | 2010 |
